# Trade-off between search costs and accuracy in oculomotor and manual search tasks

**DOI:** 10.1101/2024.10.14.618170

**Authors:** Ilja Wagner, Jan Tünnermann, Anna Schubö, Alexander C. Schütz

**Author notes:** Corresponding author: Alexander C. Schütz University of Marburg Gutenbergstraße 18, 35037 Marburg Germany.

## Abstract

Humans must weigh various factors when choosing between competing courses of action. In case of eye movements, for example, a recent study demonstrated that the human oculomotor system trades off the temporal costs of eye movements against their perceptual benefits, when choosing between competing visual search targets. Here, we compared such trade-offs between different effectors. Participants were shown search displays with targets and distractors from two stimulus sets. In each trial, they chose which target to search for, and, after finding it, discriminated a target feature. Targets differed in their search costs (how many target-similar distractors were shown) and discrimination difficulty. Participants were rewarded or penalized based on whether the target’s feature was discriminated correctly. Additionally, participants were given limited time to complete trials. Critically, they inspected search items either by eye movements only or by manual actions (tapping a stylus on a tablet). Results show that participants traded off search costs and discrimination difficulty of competing targets for both effectors, allowing them to perform close to the predictions of an ideal observer model. However, behavioral analysis and computational modelling revealed that oculomotor search performance was more strongly constrained by decision-noise (what target to choose) and sampling-noise (what information to sample during search) than manual search. We conclude that the trade-off between search costs and discrimination accuracy constitutes a general mechanism to optimize decision-making, regardless of the effector used. However, slow-paced manual actions are more robust against the detrimental influence of noise, compared to fast-paced eye movements.

**New & Noteworthy:** Humans trade off costs and perceptual benefits of eye movements for decision-making. Is this trade-off effector-specific or does it constitute a general decision-making principle? Here, we investigated this question by contrasting eye movements and manual actions (tapping a stylus on a tablet) in a search task. We found evidence for a costs-benefits trade-off in both effectors, however, eye movements were more strongly compromised by noise at different levels of decision-making.

---

Visual acuity is not uniformly high across the human eye’s retina, but it is limited to a relatively small retinal area, called the fovea. Humans use saccadic eye movements to align the fovea with different points of interest in the world, and by this, sample high-acuity visual information from relevant locations in their environment. Due to the high frequency of saccades [humans make about two saccades per second in everyday tasks (1)] and due to their low biomechanical costs, it is often believed that executing saccades as well as the corresponding gain in visual information after eye movements come at a low physiological price (cf. 2, 3). However, saccades drastically change the visual input that the brain receives and might consequently imposes a cost on visual perception.

For example, vision is impaired during ongoing saccades (4, but see 5), making the act of executing a saccade inherently costly, because it raises the risk of missing out on relevant visual information while the eye movement is in flight (6). Similarly, only one location in the environment can act as a saccade target at a time, creating additional opportunity costs due to the risk of missing out on dynamic visual information at other locations (7). Furthermore, the brain assigns costs to saccades depending on factors such as the effort it takes execute them (8–11), due to interactions between visual information in the environment and information stored in visual working memory (12–16), and due to temporal factors, such as the time it takes to find competing targets among distractors (17).

While the costs of gaze shifts seem to influence saccade execution under some conditions, this influence is not obligatory: Wagner et al. (18) showed that human observers dynamically trade off the search costs of saccades (operationalized as the time it takes to find a target stimulus among target-similar distractors) against the prospective information gain upon fixating the corresponding target (operationalized as the probability to correctly discriminate a target feature during fixation) (see also 19). This trade-off manifested in participants adapting their eye movements as well as decision-making behavior to unpredictable environmental changes in a near-optimal manner. That is, participants had an elevated probability to search for easy-to-discriminate targets, even if they carried greater search costs, and only showed an increased probability to search for difficult-to-discriminate targets when the search costs of the easy option clearly outweighed their discrimination-performance benefit. This allowed participants to perform the task close to the performance of an ideal observer model.

One question that arises from the findings by Wagner et al. (18) is whether the observed trade-off between costs and benefits of motor actions is specific for oculomotor behavior, or whether it constitutes a more general sensorimotor decision-making principle, which operates across different effectors. Some studies found striking similarities between oculomotor behavior in search tasks and search behavior when participants used manual actions instead of eye movements for information sampling (20, 21). In the paradigm by Lio et al. (20), participants either used eye movements to free-view images of naturalistic scenes, or they used manual actions (finger taps) to locally sharpen image areas that were superimposed with a Gaussian blur mask. The blur was used as a proxy for the decline in visual acuity across the retina, while the size of the area, which was sharpened after a finger tap, roughly corresponded to the size of the retinal fovea; manual actions served as proxy for eye movements as well as their visual consequences. Surprisingly, participants showed similar exploration pattern for both effectors, suggesting that similar factors were considered when choosing targets for saccadic eye movements as well as manual actions. The results by Lio et al. are in stark contrast to another line of research, demonstrating that the high biomechanical costs of manual actions and their long time course influence motor planning (22–28). As a result, participants plan sequences of manual actions further ahead compared to sequential eye movements (29).

While the study by Diamond et al. measured the isolated influence of effector-specific costs on information sampling behavior, the study by Lio et al. measured similarities and differences between information sampling behavior with different effectors. However, neither of those studies tested how effector-specific costs (e.g., longer time course and higher biomechanical costs of manual actions) are balanced against perceptual benefits of actions (e.g., information gain after eye movements or manual actions). Furthermore, neither of those studies tested whether costs and benefits of different effectors can be traded off to optimize information sampling behavior.

In the present study, we address these questions by investigating the similarities and differences between oculomotor search and manual search with a stylus and a tablet computer. More specifically, we measured decision-making behavior and tested whether the near-optimal trade-off between search costs and the information gain, previously observed for gaze shifts (18), is specific for oculomotor behavior, or whether it generalizes to manual actions. For this, we designed closely matched oculomotor and manual search tasks, allowing us to independently measure both effectors in separate tasks, and to directly compare sampling as well as decision-making behavior across different effectors.

## Methods

### Participants

We recorded data from 22 undergraduate students. One participant had to be excluded because a lack of task-understanding, resulting in a comparatively low number of valid trials (proportion valid trials of this participant: 0.60; remaining participants: *M* = 0.95, min.: 0.85, max.: 1) as well as a negative final score in the double-target condition of the oculomotor search task (final score: –0.84€; remaining participants: *M* = 3.40€, min.: 0.58€, max.: 4.98€). A second participant had to be excluded because of repeated failure to successfully calibrate the eye tracker before the double-target condition in the oculomotor search task. A third participant was excluded due to a relatively high number of gaze shifts in the single-target (proportion gaze shifts on background: 0.56; remaining participants: *M* = 0.20, min.: 0.10, max.: 0.39) and double-target condition of the oculomotor search task (proportion gaze shifts on background: 0.48; remaining participants: *M* = 0.18, min.: 0.06, max.: 0.30) that landed on the background, indicating that this participant repeatedly failed to uncover stimuli by fixating them. The remaining 19 participants had a mean age of 23.58 years (min.: 19, max.: 29 years, 11 female). Sample size was determined based on previous studies with a similar research question (16, 18, 30–32).

All participants were naïve as to the purpose of the experiment and had normal or corrected-to-normal vision. Participants were compensated with 8€/h and an additional performance-dependent bonus payout. The height of the bonus payout was determined by adding the monetary reward that participants acquired throughout conditions. The average total bonus payout across both conditions was 6.27€ (min.: 1.60€, max.: 9.72€) in the oculomotor search task, and 4.31€ (min.: 1.42€, max.: 5.58€) in the manual search task.

### Equipment

Both tasks were conducted in dimly lit lab rooms.

#### Oculomotor search task

Both conditions of the oculomotor search task were conducted using MATLAB R2016b (33) and the Psychophysics Toolbox Version 3 (34–36). A PROPixx projector (VPixx Technologies Inc., Saint-Bruno, Quebec, Canada) and a Stewart Filmscreen (Stewart Filmscreen Corporation, Torrance, California, USA) were used for stimulus presentation. The screen had a size of 90.70 × 51.00 cm, a spatial resolution of 1920 × 1080 pixel, and ran with a refresh rate of 120 Hz. The viewing distance was 106 cm. Background color was set to gray (R: 128, G: 128, B: 128, luminance: 69.6 cd/m²). The screen was calibrated to ensure linear gamma correction, and a hotspot correction was used to ensure equal luminance across the screen. Eye movements of the right eye were recorded with an EyeLink 1000+ (SR Research Ltd., Ontario, Canada), which was controlled by the Eyelink Toolbox (37). The sampling rate of the eye tracker was set to 1,000 Hz.

#### Manual search task

Both conditions of the manual search task were implemented in OpenSesame 3.3.14 (38) with custom PsychoPy (39) presentation routines. Conditions of the manual search task were presented on a WACOM MobileStudio Pro 16 (Kazo, Saitama, Japan) tablet-PC with a 345 × 195 mm stylus/touch display, running at a spatial resolution of 1280 × 720 pixels and a 60 Hz refresh rate. The tablet was tilted by 32° to achieve more planar viewing. The viewing distance varied based on participant height, and varied approximately between 60 and 90 cm. The stimulus display background color (R: 128, G: 128, B: 128) had a luminance of 34.53 cd/m² on the tablet screen.No eye movements were recorded for the manual search task.

### Stimuli

In all tasks and conditions, we used the same stimuli as Wagner et al. (18). A combination of cross and bull’s eye [total diameter: 0.60 degrees of visual angle (dva)] was used as fixation cross (40). Custom stimuli, made out of a colored circular outline and a round-edged rectangle were used as targets, distractors, and mask stimuli (Figure 1A). The circular outline was either red (R: 255, G: 0, B: 0; luminance projector screen: 43.70 cd/m²; luminance tablet screen: 44.76 cd/m²) or blue (R: 0, G: 0, B: 255; luminance projector: 6.10 cd/m²; luminance tablet:17.36 cd/m²). It had a diameter of 1.20 dva, and a line thickness of 0.10 dva. The rectangle was gray (R: 109, G: 109, B: 109; luminance projector: 59.70 cd/m²; luminance tablet: 27.08 cd/m²), had a line thickness of 0.03 dva, and it’s longer and shorter side had a length of 0.69 dva and 0.23 dva, respectively. Although the absolute values differed between the projector and the tablet setup for the oculomotor and the manual search, respectively, the Weber contrast of the task-relevant rectangles relative to the background was fairly similar (projector: 14 %; tablet: 21 %). Targets were defined by the rectangular part of stimuli either being oriented horizontally or vertically, while inner rectangles of distractors were oriented oblique (45° or 135°). Mask stimuli were generated by superimposing rectangles of all orientations onto each other (Figure 1A).

**Figure 1.**
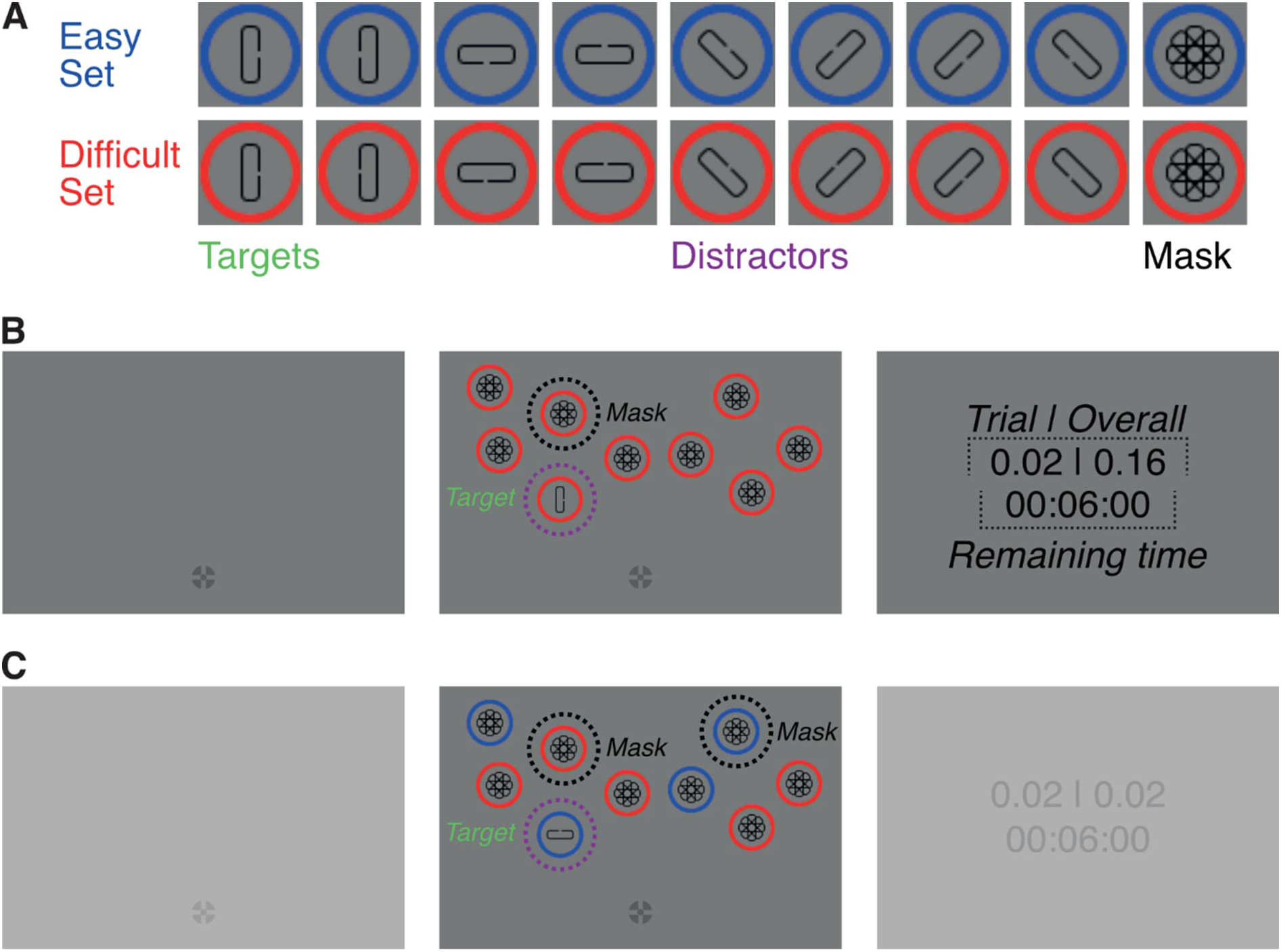
Stimuli and trial procedure. (A) Stimuli were oriented rectangles, centered within blue or red circles. Based on the circle color, stimuli formed two sets: the easy and difficult set. Each set consisted of one target, several distractor variations, and a mask stimulus. Stimuli from the two sets differed in respect to the size of a gap, located on one side of their rectangles, so that the gap position was more difficult to discriminate on stimuli from one set (easy set) than the other (difficult set). (B) Participants in the single-target condition were instructed to find a target, placed among a variable number of distractors (here: eight), and to report the gap position on the target. In each trial, only elements from one set (here: the difficult set) were shown. (C) Participants in the double-target condition had the same task as in the single-target condition. However, elements from both sets were shown in each trial of the double-target condition. (B)–(C) Participants received feedback about their response and the time remaining to complete trials after each trial. Stimuli were masked, unless participants gaze (oculomotor search task) or the stylus (manual search task) were located within an area of interest around a stimulus. Differences between conditions are highlighted by increasing the brightness of screens that did not change; this only applies to this illustration and was not part of the actual experiment Stimuli are not drawn to scale. Dashed lines/circles and italic text were not part of the display and are shown for illustration purposes only. (A)–(C) For illustration purposes, rectangles within circles are drawn in black, whereas they were drawn in gray during the task.

Stimuli were divided into two sets: the easy and difficult set. Set affiliation was based on the color of the circular outline of stimuli. Stimuli of the easy set had a comparatively large gap on one of the long sides of their respective rectangle (gap size: 0.08 dva), while the gap size on stimuli from the difficult set was comparatively small (0.05 dva). Gap sizes were chosen based on values reported by Wagner et al. (18), such that participants had a higher probability of correctly reporting the gap position on stimuli from the easy set. Mask stimuli had no gap. Anti-aliasing was applied to circular outlines as well as rectangles. For targets, anti-aliasing only occurred at the round edges of rectangles but not to the rectangle’s long sides. This was done to avoid interferences between task-performance and blurring due to anti-aliasing.

In each trial, stimuli were randomly distributed across a screen area, measuring 16 dva × 16 dva. For this, we drew random coordinates from within the designated area until each stimulus in a trial was placed. While drawing coordinates, it was ensured that an equal number of stimuli were shown in the left and right as well as in the upper and lower half of the stimulus area. Additionally, it was ensured that the Euclidean distance between all stimuli, including the fixation cross, was at least 4 dva. This was done to ensure that participants had to fixate individual stimuli to discriminate targets from distractors, and that they could not use peripheral vision to do so. The lower end of the stimulus area was located 4 dva and the upper end 20 dva above the location of the fixation cross. The stimulus area extended 8 dva to the left and right of the vertical screen center. The fixation cross was placed at the horizontal screen center, and at an off-center vertical location, 9.5 dva beneath the vertical screen center.

Note that in the manual search tasks the distance between display and participant eyes – and consequently the apparent stimulus size – was variable, depending on participant body size (not measured), to allow for natural and unobstructed interaction with the tablet. Stimulus placement and sizes were selected to match the values reported above at a typical viewing distance of 61 cm.

### Design

We used a within-subject design. Each participant was asked to complete two tasks the manual and oculomotor search task. The tasks were completed in a single session on a single day. The order, in which tasks were completed, was counterbalanced, so that half of the participants first completed the oculomotor search task, while the other half of participants began with the manual search task. In each task, participants completed two conditions: the single- and double-target condition. Both conditions were completed sequentially, and participants always began with the single-target condition.

Stimuli of the easy and difficult set were indicated by blue/red circles for even participant numbers and vice versa for odd participant numbers. We counterbalanced the color of the circular outlines around stimuli, so that the circle around stimuli from the easy set was red for participants with even participant numbers. This assignment was inverted for participants with odd participant numbers. The individual mapping between outline color and set affiliation was kept constant across tasks and conditions, for each participant. The orientation of target stimuli was randomized across trials. When both targets were shown together in a trial, it was ensured that both targets were shown with different orientations.

Participants in each condition of both tasks were given 6 min and 30 s to complete as many trials as they can. To ensure that we collect an approximately equal number of trials for each combination of target difficulty and number of same-colored distractors, trials in the single-target condition were presented as miniblocks. Here, one miniblock consisted of 18 trials (one out of two targets presented together with 0–8 same-colored distractors), while the order of individual trials within a miniblock was randomized. Similarly, trials in the double-target condition were also presented as miniblocks. However, here, one miniblock consisted of nine trials (both targets presented together with eight distractors, from which 0–8 were drawn from the easy set and 8–0 were drawn from the difficult set).

Participants completed 36 practice trials before each condition (i.e., each participant completed 4 × 36 practice trials in total). This relatively large number of demonstration trials was included to ensure that participants were well familiar with the respective setup before running the actual measurement. Participants were given the opportunity to rest between tasks as well as between conditions.

### Procedure

We used the same procedure as Wagner et al. for oculomotor search (18). Before each condition, participants were instructed about what task they have to perform (see below), whether one or two targets will be presented in trials, how the two targets differed regarding gap size on the rectangular inner element, and about the relationship between gap size and the color of the outer circle. Additionally, participants were instructed that each stimulus in a trial can only be viewed for a limited total duration after which it would remain masked for the remainder of the trial.

For the manual search task, participants were additionally instructed to use the tablet stylus with their dominant hand. They were asked to hold the stylus like a normal pen but, if needed, to hold it more towards its end, so that they could easily reach stimuli everywhere on the tablet. They were instructed to touch the outer circle of stimuli with the stylus tip in a position so that they would not occlude the stimulus with their hand. Participants were told that the stylus had to be lifted and not dragged between stimuli, when moving from one stimulus to the next.

#### Single-target condition

The single-target condition served two purposes. First, it was intended to familiarize participants with the stimuli as well as the general task. Second, we used the single-target condition to quantify, for each target individually, the accuracy by which participants could detect the gap on targets as well as various temporal measures of search performance (see Data Analysis).

Participants started the trials with their dominant hand. In the oculomotor search task, participant looked at the fixation cross and either pressed the spacebar (left-handed participants) or the Enter-key on the numpad (right-handed participants) (Figure 1B). In the manual search task, participants had to touch the fixation cross with the stylus and hold it down until stimulus display onset to initiate the trial. Stimulus display onset occurred after a random interval, drawn from a uniform distribution with a lower limit of 500 ms and an upper limit of 1,000 ms.

In each trial, the stimulus display consisted of one target and a variable number of distractors (0–8) from the same set as the target. Participants were instructed to find the target and report the location of the gap by pressing either the up or down arrow key (for horizontal targets where the gap was on the upper or lower edge) or the left or right arrow key (vertical targets) (Figure 1A). To minimize accidental responses, the active response keys in a trial were, however, confined to the orientation of the target that was shown in a trial, e.g., if the target in a trial was oriented horizontally, only the up- and down-key were active.

To make the oculomotor and manual search tasks as comparable as possible, stimuli were masked, and participants had to either bring their gaze (oculomotor search task) or the stylus (manual search task) within an AOI around a stimulus to reveal if it was a target or distractor (see Online tracking of inspected AOI). Trials were untimed and participants could take as much time as they want to complete trials. Each stimulus in a trial, however, could only be viewed for up to 500 ms, before it was masked for the remainder of the trial. The stimuli (including the fixation cross) stayed on until participants provided a response.

After a response was detected, the stimuli disappeared, and feedback was shown at the center of the screen (Figure 1B, right panel). The feedback informed participants about how much time they had left to complete trials, their current score, and whether their most recent response was correct or not. Participants received two points for correct responses and lost two points for incorrect ones. The remaining time to complete trials was updated after each trial by subtracting the time elapsed between onset of the stimulus display and response. In the oculomotor search task, a red error message (“Not fixated!”) was shown for 1,500 ms instead of the performance feedback if participants moved their gaze more than 1.50 dva from the fixation cross before onset of the stimulus display. Whenever such an error occurred, two points were subtracted from the participant’s score, irrespective of whether the response in a trial was correct, and the remaining time was updated in the same way as in trials without fixation error. No such proximity check was implemented in the manual search experiment.

#### Double-target condition

The double-target condition was used to measure how participants balanced search costs and accuracy when choosing between competing target options. It was mostly identical to the single-target condition, and only differed in regard to details of the task and the composition of stimulus displays (Figure 1C). First, each stimulus display contained two targets, with one target belonging to the easy set, and the other being part of the difficult set. Participants were instructed that they can freely choose which of the two targets to report. Second, stimulus displays in the double-target condition always contained eight distractors, from which 0–8 distractors were drawn from the easy, and 8–0 distractors were drawn from the difficult set. Thus, stimulus displays in the double-target condition always contained ten elements in total, from which 1–9 elements belonged to the easy set and 9–1 element belonged to the difficult set. However, the exact composition of stimulus displays was randomized over trials. Third, since both targets were shown in each trial of the double-target condition, all response keys were active in each trial.

The paradigm, used in the double-target condition, has some notable similarities but also important differences to a paradigm recently introduced by Irons & Leber (17, 41). Those similarities include the pairing of a search with a decision-making task as well as the presentation of targets with varying search costs. The differences include the additional manipulation of discrimination difficulty of the two targets and the random variation of search costs across trials. For a more in-depth discussion of the similarities and differences between the two paradigms we would like to refer the reader to the corresponding comparison in the Methods section of Wagner et al. (18).

### Data analysis

#### Online tracking of inspected AOIs

Stimuli in both tasks were masked unless they were uncovered by the participant’s gaze (oculomotor search task) or the stylus (manual search task). Gaze had to be located within an AOI around a stimulus (diameter: 3 dva, relative to stimulus center; invisible to participants), whereas the stylus needed to be on a ring with 0.76 dva inner and 1.47 outer diameter, to motivate clicking the ring without making it too difficult. Whenever gaze or the stylus were detected to be within an AOI, the corresponding stimulus was unmasked, revealing whether it was a target or distractor. When gaze or the stylus left an AOI, the corresponding stimulus was masked again. For the oculomotor task, stimuli were masked with a 250 ms delay after the gaze left the corresponding AOI. This was done to avoid flickering of stimuli when participants gaze dwelled on the border of an AOI, and by this, drifted in and out of it. In the manual search task, where there is less jitter in trajectories, the stimuli were masked without delay. Whenever the eye tracker detected a signal loss (e.g., during a blink), each unmasked stimulus was masked, until gaze was again detected to be within an AOI.

Additionally, each stimulus in a trial could be viewed for a total of 500 ms (that could also be spent across several visits of the stimulus) before it was replaced by a mask for the remainder of a trial. In the oculomotor search task, the viewing time of individual stimuli was tracked by determining when a participant’s gaze entered as well as left an AOI, and by subtracting the difference between those two timestamps from the overall permitted viewing time of this stimulus. Although our tracking algorithm could determine when gaze entered and left an AOI, it was agnostic to whether gaze was stationary during an AOI visit. Consequently, instances where gaze was stationary and dwelled within an AOI as well as instances where gaze merely passed through an AOI midflight (e.g., during saccades) were both treated as AOI visits, resulting in an update of the overall permitted viewing time. A similar procedure was used to track stimulus viewing in the manual search task: The viewing time was determined as the difference between the time the stylus uncovered the stimulus and when it was masked on stylus exit.

#### Eye movement analysis

On- and offsets of saccades and blinks were detected using the EyeLink algorithm. Since saccades and blinks can both move the eye, and consequently change which AOI the gaze is currently located in, both types of eye movements were analyzed together. Visited AOIs were determined offline by using the same logic as for the online detection (see Online tracking of inspected AOI). To reduce the influence of noise, we used the average gaze position between the offset and onset of neighboring gaze shifts to determine whether gaze was located within an AOI. Repeated visits of the same AOI were treated as independent instances and analyzed separately. Thus, our analysis does not account for factors such as the integration/accumulation of information across multiple visits of the same stimulus.

We excluded gaze shifts from analysis that had durations shorter than 5 ms, whose offsets occurred after stimulus array offset, and whose onset/offset coordinates lay outside of screen bounds. Additionally, we excluded corrective gaze shifts that were made within AOIs, i.e., small gaze shifts that did not change the currently inspected AOI, as well as the last gaze shift in a trial (i.e., the gaze shift immediately before a response) if the last gaze shift targeted anything but an AOI (e.g., the screen background).

#### Analysis of manual actions

On- and offset of the manual movements were obtained from the timestamps of stimulus uncovering and masking events logged by the experiment.

#### Trial exclusion

In the oculomotor search task, we excluded trials from analysis when participant’s gaze, at any time within a window of −20–80 ms around onset of the stimulus display, deviated more than 1.50 dva from fixation. Additionally, some trials were excluded due to technical difficulties during data recording.

In the manual search task, we excluded trials where participants dragged the stylus across the screen, instead of lifting it when moving from one stimulus to the next. Even though participants were instructed to avoid pen dragging, one could consider it a valid strategy to explore the displays. However, pen dragging interfered with reliably detecting movement on- and offsets. Hence, we checked the trials against three criteria that indicate pen dragging and removed the affected trials from the analysis. The criteria and the detection procedure are described in Appendix X.

After trial exclusion in the oculomotor search task, we were left, on average, with 95.10% (min.: 84.62%, max.: 99.59%) valid trials in the respective single-target condition, and 95% (min.: 84.50%, max.: 99.63%) valid trials in the respective double-target condition. After trial exclusion in the manual search task, we were left, on average, with 97.93% (min.: 81.48%, max.:100%) valid trials in the respective single-target condition, and 98.90% (min.: 89.68%, max.:100%) valid trials in the respective double-target condition.

#### Calculating planning, inspection, and response time in the oculomotor search task

In the oculomotor search task, planning times were calculated as the difference between the time of stimulus display onset and offset of the first gaze shift with onset after stimulus onset. Response times were calculated as the difference between the time of response and offset of the last gaze shift before a response. Inspection times were calculated by, first, determining which stimuli were fixated in a trial, second, calculating the time participants spent dwelling within the AOI of each fixated stimulus, and third, calculating the median of the resulting vector to obtain an estimate for the average inspection time in a trial.

To account for differences in information uptake, dwell times of individual stimuli were calculated slightly differently depending on whether an AOI was left by means of a saccade or a blink. If an AOI was left via a saccade, dwell times were determined by calculating how much time elapsed between offset of the gaze shift that brought gaze to an AOI and offset of the saccade that resulted in the gaze leaving the AOI. If an AOI was left via a blink, the corresponding dwell time was calculated using the difference between the timestamp of the gaze shift that brought gaze to the AOI and onset of the blink which caused gaze to leave the AOI. If a blink occurred while gaze was located within an AOI (i.e., the gaze position was within the same AOI before and after the blink), the duration of the blink was subtracted from the corresponding dwell time.

No inspection times were calculated in the single-target condition, whenever gaze was located within the AOI around a target, as this constituted the last fixated element before the response. In the double-target condition, inspection times were calculated irrespective of whether gaze was located within the AOI around target or a distractor. The only exception to this were instances where gaze was located within the AOI around a target while a response was given (i.e., the last inspected stimulus before a response was a target). If participants responded during an ongoing gaze shift the corresponding gaze shift was ignored, and the corresponding case treated as if the target was fixated until response. The same logic was applied to cases where participants gave a response while their gaze was located outside any AOI (e.g., on the screen background), immediately after fixating a target. For planning, inspection, and response times, outliers were removed from the corresponding data distributions. Outlier were detected on a single-participant level, using MATLAB’s isoutlier function. A trial was flagged as outlier, whenever the corresponding temporal measure deviated more than three scaled median absolute deviations from the median.

#### Calculating planning, inspection, and response time in the manual search task

The same processing steps were applied to temporal measures in the manual search task as to temporal measures in the oculomotor search task.

#### Statistical analysis

Two-by-two repeated measures ANOVAs were used for statistical inference. Additionally, since data distributions were often skewed and contained outlier, we opted to use Yuen’s modified *t*-test for trimmed dependent means (42) as a robust alternative to the conventional *t*-test. For robust *t*-tests, we used the recommended trimming of 20% (43). We used Spearman’s rho to calculate correlation coefficients for the same reason. The robust *t*-test was implemented as part of the WRS2 package for R (44). ANOVAs were run using JASP 0.19 (45).

For proportion gaze shifts and accuracy as well as planning-, inspection- and response-times we, first, calculated the corresponding variable for each set size condition individually, and second, averaged over the resulting vector to obtain the final variable estimate for a participant. Since distributions of planning-, inspection-, and response-times tended to be skewed, we used the median to aggregate over set-size-conditions, while we used the mean for all other variables. Additionally, we applied an arcsine-transformation to proportion data to account for the fact that the underlying data distribution was highly skewed due to a large number of values close to one, especially in the manual search condition (e.g., Figure 2A&E). For consistency, all ANOVAs with proportional data as dependent variable were run on arcsine-transformed data; the values, reported and plotted throughout this manuscript, are untransformed values.

**Figure 2.**
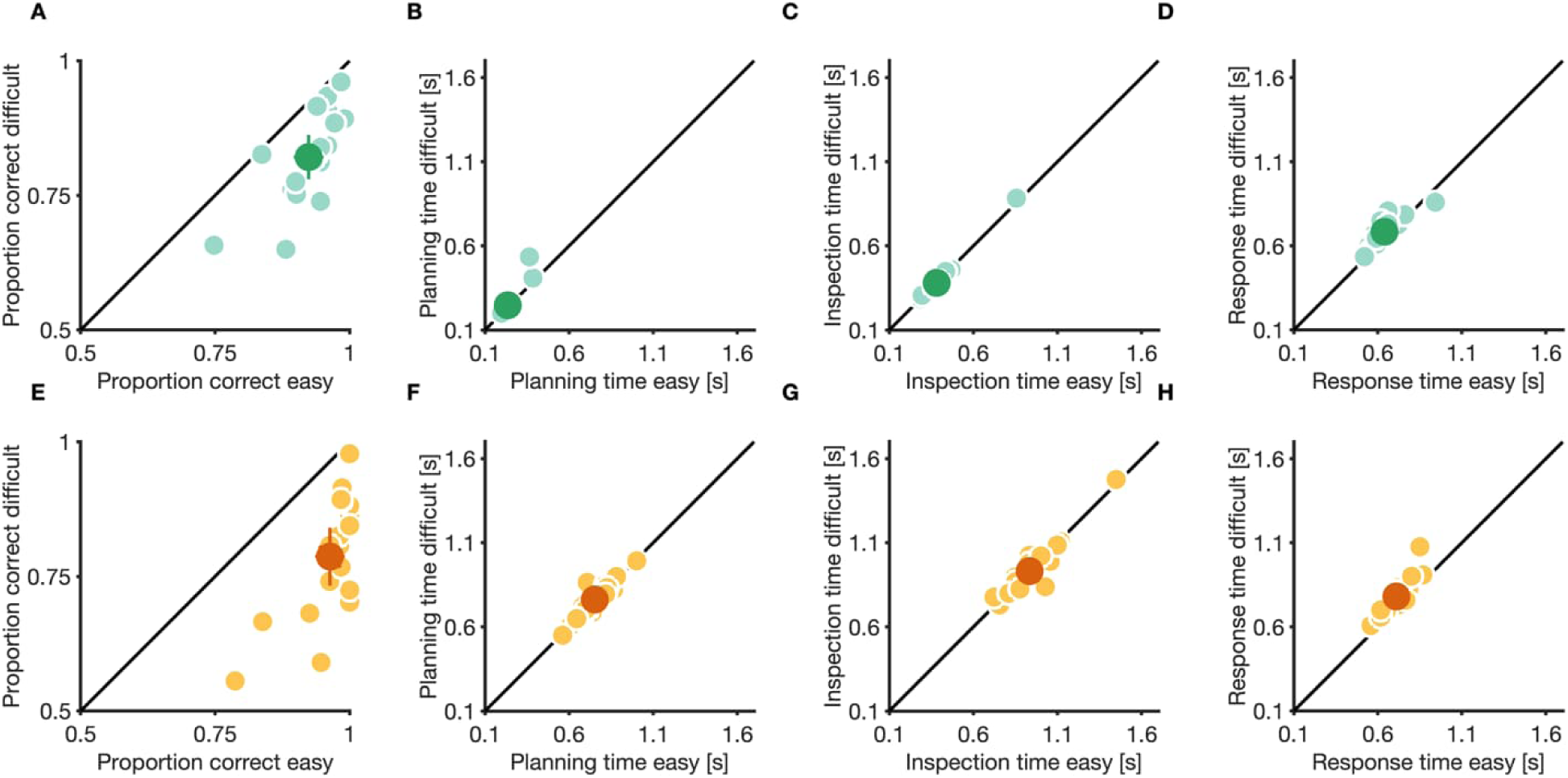
Perceptual performance and search behavior in the single-target condition. (A&E) Discrimination performance, (B&F) planning time, (C&G) inspection time, and (D&H) response time for easy and difficult targets of the oculomotor search (A–D) and manual search task (E–H). Small, light dots are data from individual participants, large, dark dots are means. Error bars are 95% confidence intervals.

Unlike in our previous publication (18), where we used linear models to estimate the contribution of search costs and discrimination difficult to choice behavior, we fitted sigmoids (Gaussian cumulative density function) to data in this study. This was done because linear models, especially in the manual search task, often provided a relatively poor fit to the data. To avoid unreasonable parameter estimates, we restricted the mean parameter of sigmoids to be in the range of [-2, 10], while the standard deviation parameter was unconstrained. The range of the mean of sigmoids was chosen to reflect the range of set-size-conditions, as shown in the experiment. Since standard deviations of sigmoids were treated as free parameter and had no constraints, they could theoretically assume any value, including negative ones. A positive standard deviation corresponds to a positive relationship of easy set size and proportion of easy choices, a negative standard deviation corresponds to a negative relationship between easy set size and proportion of easy choices. To estimate a slope (similar to the linear function in our previous study), we calculated the Gaussian probability density (as the first derivative of the cumulative Gaussian function) at a mean of zero and the fitted standard deviation of the cumulative Gaussian function. Whenever the estimated standard deviation was positive, we multiplied the calculated slope by -1.

### Modeling

We used the same generative stochastic model as Wagner et al. (18) to predict participant’s choice and sampling behavior in the double-target conditions of both tasks. In short, the model assumes that participants, in general, act like an ideal observer, who has full knowledge about its own performance limitation, who can accurately estimate the relative gain of competing target options, and who aims to maximize its accumulated reward by consistently choosing to search and discriminate higher-gain targets. However, the gain estimates of the ideal observer are corrupted by a variable amount of random decision-noise, leading to more or less prevalent choices for lower-gain targets. Additionally, the model assumes that the ideal observer will choose a target at trial start, before starting search, and subsequently only inspect elements from the set of the chosen target until the chosen target is found. This preference for sampling elements from the chosen set (sampling bias) is, however, corrupted by additional, independent random sampling-noise, which can lead to occasional sampling of elements from the set of the non-chosen target.

We implemented this general logic in two steps: First, we used data from the single-target condition of either the oculomotor or manual search task to generate ideal observer predictions about the relative gain of available target options for each possible relative number of easy and difficult distractors on the screen. For this, we used data about participants discrimination accuracy and various measures of their temporal search performance, i.e., individual planning, inspection, and response times. Those gain estimates provided a normative guideline for how a participant, who strives to maximize their accumulated reward, should optimally act, given a relative number of easy and difficult distractors on the screen. Those ideal observer predictions were, in a second step, used as input to the generative stochastic model, where they were corrupted by decision- as well as sampling-noise, and where they were used to calculate a participant’s individual target preference and sampling bias. The strength of both sources of noise was controlled by two free model parameters. Critically, the model only received gain estimates as input and ignored external factors, such as the placement of stimuli on the screen as well as the spatial distance between stimuli, and internal factors, such as differences in the metabolic costs of the effector used to search for targets.

The model returned two predictions. First, the model returned a vector with probabilities that a participant with a given decision-noise parameter will choose to discriminate easy targets, given a relative number of easy and difficult distractors on the screen. Second, the model returned another vector with probabilities that a participant with a given sampling-noise parameter will sample elements from the set of the chosen target while searching for the target, again, given a relative number of easy and difficult distractors on the screen. To avoid unreasonable values for free parameter, the parameter bounds were set as [0, 2.50] for both, sampling- and decision noise. For a more extensive and formal description of the generative stochastic model we would like to refer the reader to the original publication by Wagner et al. (18), where we first introduced this modelling framework.

## Results

To test whether the previously observed, near-optimal trade-off between search costs and discrimination accuracy in a perceptual task is specific for gaze shifts, or whether it generalizes to manual actions, we independently measured gaze shifts and manual actions in two tasks (oculomotor and manual search task). Although both effectors were measured independently, we designed the tasks to be as similar as possible, allowing us to directly compare sampling and choice behavior between effectors. Each task consisted of two conditions (single- and double-target), each serving a different purpose. While the single-target condition was used to familiarize participants with the paradigm and assess their target-specific search performance in isolation, the double-target condition was used to test for trade-offs between search costs and discrimination difficulty of competing target options.

### Manual search carries a larger temporal cost than oculomotor search

To assess participant’s target-specific search performance in the single-target conditions of the two tasks, we measured their discrimination performance and various temporal measures. More specifically, temporal measures included planning time (average time between stimulus onset and offset of first movement), inspection time (average dwell time on stimuli in trials), and response time (time between offset of last movement before response and time of response). Each measure was quantified separately for each target and each task. Since we explicitly manipulated discrimination difficulty of targets, we expect higher accuracy for easy than for difficult targets, but no differences in other measures of search performance. We hypothesize that this pattern holds for both, the oculomotor and the manual search task.

In the single-target condition of the visual search experiment, participants had a higher discrimination performance for easy (proportion correct *M* = 0.92, *CI*_95%_ [0.90, 0.95]) compared to difficult targets (*M* = 0.82, *CI*_95%_ [0.78, 0.86]). Furthermore, response times were also lower for easy (*M* = 639 ms, *CI*_95%_ [592 ms, 686 ms]) compared to difficult targets (*M* = 684 ms, *CI*_95%_ [642 ms, 725 ms]). However, neither planning times (easy target: *M* = 237 ms, *CI*_95%_ [212 ms, 261 ms]; difficult target: *M* = 247 ms, *CI*_95%_ [207 ms, 288 ms]), nor inspection times (easy target: *M* = 382 ms, *CI*_95%_ [322 ms, 443 ms]; difficult target: *M* = 380 ms, *CI*_95%_ [317 ms, 444 ms]), differed substantially between target options in the single-target condition of the oculomotor search task.

In the single-target condition of the manual search task, participants showed a similar pattern in their search behavior: their discrimination performance was higher for easy (*M* = 0.96, *CI*_95%_ [0.94, 0.99]) compared to difficult targets (*M* = 0.79, *CI*_95%_ [0.73, 0.84]), and their response times where lower for easy (*M* = 710 ms, *CI*_95%_ [672 ms, 749 ms]) compared to difficult targets (*M* = 780 ms, *CI*_95%_ [728 ms, 831 ms]). Again, neither planning times (easy target: *M* = 754 ms, *CI*_95%_ [703 ms, 804 ms]; difficult target: *M* = 762 ms, *CI*_95%_ [710 ms, 814 ms]), nor inspection times (easy target: *M* = 934 ms, *CI*_95%_ [851 ms, 1,017 ms]; difficult target: *M* = 932 ms, *CI*_95%_ [848 ms, 1,015 ms]), differed substantially between target options.

In line with this pattern, a 2-by-2 repeated measures ANOVA with task as well as target as factors and perceptual performance as dependent variable showed no significant main effect of task, *F*(1,18) = 3.02, *p* = 0.100, but a significant main effect of target, *F*(1,18) = 183.83, *p* < 0.001, as well as a significant interaction between target and task, *F*(1,18) = 32.07, *p* < 0.001. For planning times, the ANOVA showed a significant main effect of task, *F*(1,18) = 451.61, *p* < 0.001, but neither a main effect of target, *F*(1,18) = 2.12, *p* = 0.156, nor an interaction between the factors, *F*(1,18) = 0.026, *p* = 0.874. Similarly, for inspection times, the ANOVA showed a significant main effect of task, *F*(1,18) = 504.10, *p* < 0.001, but neither a main effect of target, *F*(1,18) = 0.094, *p* = 0.762, nor an interaction between the factors, *F*(1,18) = 0.0002, *p* = 0.989. However, for response times, the ANOVA showed significant main effects for both target, *F*(1,18) = 41.88, *p* < 0.001, and task, *F*(1,18) = 14.42, *p* = 0.001, but no interaction between the factors, *F*(1,18) = 2.54, *p* = 0.129.

The higher temporal expenditure in the manual search task was reflected in participant’s performance: in the manual search task, participants completed fewer trials (*M* = 115.05 trials; min.: 67 trials, max.: 171 trials) and received a lower overall bonus payout (*M* = 1.69€; min.: 0.46€, max.: 2.50€), compared to the average number of completed trials (*M* = 209.68 trials; min.: 104 trials, max.: 261 trials), *Y_t_*(12) = 14.37, *p* < 0.001, and the average bonus payout in the oculomotor search task (*M* = 2.88€; min.: 0.44€, max.: 4.74€), *Y_t_*(12) = 8.74, *p* < 0.001.

In summary, the different gap sizes of easy and difficult targets influenced participant’s perceptual discrimination performance and their response times, while the same manipulation did not impact other measures of search behavior, neither in the oculomotor nor the manual search task. However, although the general pattern in participant’s search behavior was similar in the single-target conditions of the two tasks, the higher temporal costs of manual search negatively impacted participant’s search performance in this task.

### Participants trade off search costs and discrimination difficulty for decision-making irrespective of effector

While in the single-target conditions of both tasks, participants saw one target (easy or difficult) as well as a variable number of distractors from the same set as the target, we presented both targets as well as a mix of both distractor types in trials of the double-target conditions. Thus, in the double-target conditions of both tasks, participants not only had to find and discriminate a target, but they also had to make decisions about which target to search and discriminate.

Generally, in the double-target conditions, participants completed more trials than in the respective single-target condition of the oculomotor (*M* = 227.89 trials; min.: 95 trials, max.: 277 trials), *Y_t_*(12) = -5.00, *p* < 0.001, and manual search task (*M* = 152.68 trials; min.: 84 trials, max.: 190 trials), *Y_t_*(12) = -14.50, *p* < 0.001. Similarly, in the double-target conditions, participants accumulated a higher bonus payout than in the respective single-target condition of the oculomotor (*M* = 3.40€; min.: 0.58€, max.: 4.98€), *Y_t_*(12) = -4.36, *p* < 0.001, and manual search task (*M* = 2.62€; min.: 0.96€, max.: 3.34€), *Y_t_*(12) = -11.96, *p* < 0.001.

One reason for this performance benefit might be that participants in the respective double-target condition were aware of their own performance limitations (probability to correctly discriminate each target option, time needed to find a target given the number of same-colored distractors on the screen) and used this knowledge to optimize decision-making behavior. In other words, in the double-target conditions of both tasks, participants might have acted like a theoretical ideal observer who considers the relative search costs and discrimination difficult of available target options, and always chooses targets with higher expected value (i.e., higher gain over time). To test whether participants considered relative search costs and discrimination difficulty for decision-making, we fitted sigmoid functions (Gaussian cumulative density functions) to the choice data of participants (Figure 3A–B) and analyzed the means and slopes of the resulting fits (Figure 3C).

**Figure 3.**
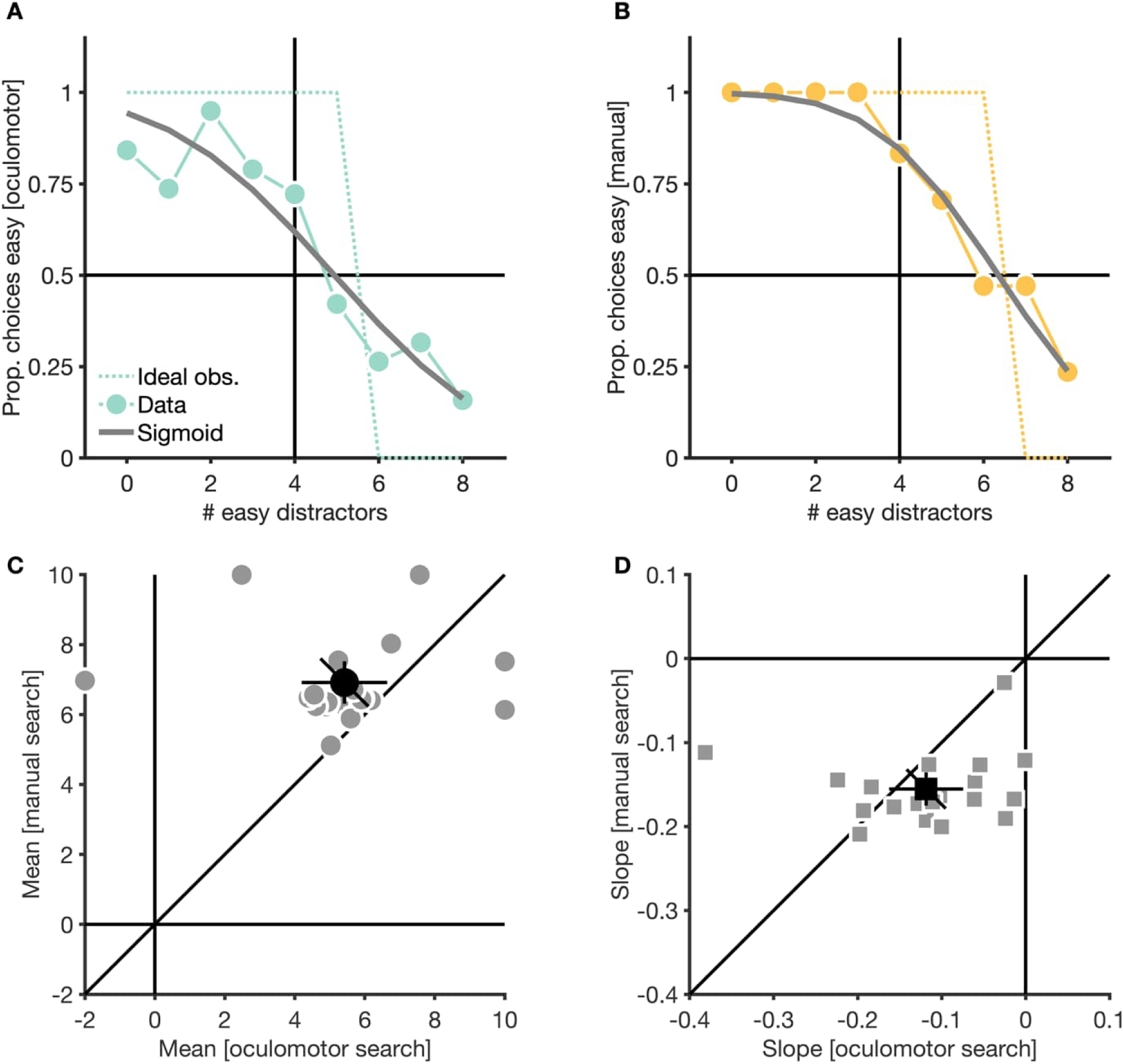
Influence of search costs and discrimination difficulty on choice behavior. (A–B) Proportion of trials in which one representative participant in the double-target condition of the oculomotor (A) and manual search task (B) chose to discriminate easy targets. Small, light dots are data, gray solid lines are fitted sigmoid functions, light dotted lines are ideal observer predictions. (C)–(D) Means (C) and slopes of sigmoid functions (D), fitted to the choice data of participants in the double-target conditions of the two tasks. Small, light dots are data from individual participants, larger, black dots are means. Error bars are 95% confidence intervals.

Aggregated means of sigmoids were smaller in the oculomotor (*M* = 5.42; *CI*_95%_ [4.19, 6.65]) than in the manual search task (*M* = 6.92; *CI*_95%_ [6.31, 7.52]), *Y_t_*(12) = -4.18, *p* = 0.001, suggesting that participants had a more pronounced preference for easy targets in the manual search task, which persisted even when the search costs of the easy and difficult set where equal. Slopes of sigmoids, on the other hand, where larger in the oculomotor (*M* = - 0.12; CI_95%_ [-0.16, -0.07]) than the manual search task (*M* = -0.16; *CI*_95%_ [-0.18, -0.14]), suggesting steeper sigmoid functions in the manual search task, and thus, a greater weighting of search costs in this task, *Y_t_*(12) = 2.41, *p* = 0.033. Interestingly, although participants did consider the relative search costs and discrimination difficulty of targets for decision-making, they did not consistently choose like an ideal observer would choose. Instead, participants occasionally chose lower gain targets in both the oculomotor search (Figure 3A & Figure S1) and manual search task (Figure 3B & Figure S2).

In summary, we found that in the double-target condition of both tasks, participants optimized their decision-making behavior by trading off the relative search costs and discrimination difficulty of target options. In line with the greater temporal costs of manual compared to oculomotor search (Figure 2B–D&F–H), search costs where weighted stronger in the manual search task. However, we also observed a stronger bias to search for easy targets in the manual search task, even if searching for this target carried larger search costs. This might be due to the somewhat larger difference in discrimination difficulty between the easy and the difficult target in the manual compared to the oculomotor search task.

### Visual information sampling behavior is strongly driven by heuristics

Besides target choices, participants’ performances in our task also depended on their information sampling behavior during search for targets. An ideal observer should make the decision about which target to search for immediately after onset of the stimulus display and delay or suppress any gaze shifts (oculomotor search) as well as manual actions (manual search) until a decision was made. Suppressing such premature movements will allow an ideal observer to sample information exclusively from the set of the chosen target (as opposed to random sampling or choosing a target while already sampling information from the search display), and not waste time on sampling information that is not instrumental for finding the chosen target. To quantify participants’ information sampling behavior during search, we analyzed the probability of individual movements in a trial to target elements from the set of the eventually chosen target. We focused on the first two movements in trials, due to large differences in the overall number of movements made between participants.

In the oculomotor search task, information sampling during search was initially unbiased, with participants having a close to equal probability for sampling elements from the set of the chosen and not-chosen target (*M* = 0.54; *CI*_95%_ [0.48, 0.59]). Only with the second gaze shift participants showed a clear bias towards elements from the set of the eventually chosen target (*M* = 0.72; *CI*_95%_ [0.66, 0.77]). Gaze shifts, furthermore, showed general biases for elements from the easy set, the smaller set as well as elements closest to the current fixation location. The bias for easy elements was comparable between first (*M* = 0.53; *CI*_95%_ [0.48, 0.58]) and second gaze shifts (*M* = 0.61; *CI*_95%_ [0.55, 0.66]), while the bias for elements from the smaller set was less pronounced for first gaze shifts (*M* = 0.37; *CI*_95%_ [0.32, 0.42]) than for second gaze shifts (*M* = 0.54; *CI*_95%_ [0.46, 0.63]). The bias for close elements, on the other hand, was most pronounced for first gaze shifts (*M* = 0.82; *CI*_95%_ [0.77, 0.88]), and declined strongly for the subsequent gaze shift, (*M* = 0.42; *CI*_95%_ [0.40, 0.45]). In line with this general pattern, we observed shorter latencies for first (*M* = 173 ms; *CI*_95%_ [142 ms, 203 ms]) than for second gaze shifts (*M* = 208 ms; *CI*_95%_ [171 ms, 245 ms]).

In the manual search task, information sampling during search was strongly biased towards elements from the set of the eventually chosen target, and the strength of this bias was similar for the first (*M* = 0.84; *CI*_95%_ [0.81, 0.87]) and second stylus movement in a trial (*M* = 0.85; *CI*_95%_ [0.81, 0.88]). Similar to the oculomotor search task, information sampling in the manual search task was also biased towards elements from the easy set, the smaller set as well as elements closest to the current location of the stylus. However, all biases were similar for the first (easy-set-bias: *M* = 0.63; *CI*_95%_ [0.58, 0.69]; smaller-set-bias: *M* = 0.84; *CI*_95%_ [0.78, 0.90]; proximity bias: *M* = 0.22; *CI*_95%_ [0.18, 0.25]) and second movement in a trial (easy-set-bias: *M* = 0.65; *CI*_95%_ [0.59, 0.71]; smaller-set-bias: *M* = 0.79; *CI*_95%_ [0.71, 0.88]; proximity bias: *M* = 0.16; *CI*_95%_ [0.12, 0.19]). Similar to the oculomotor search task, however, latencies of movements in the manual search task were shorter for first (*M* = 317 ms; *CI*_95%_ [293 ms, 341 ms]) than for second movements (*M* = 411 ms; *CI*_95%_ [380 ms, 443 ms]).

In line with this pattern, a 2-by-2 repeated measures ANOVA with movement and task as factors and proportion movements on the set of the chosen target as dependent variable yielded a significant main effect for movement, *F*(1,18) = 29.17, *p* < 0.001, a significant main effect for task, *F*(1,18) = 73.47, *p* < 0.001, and a significant interaction between those factors, *F*(1,18) = 31.50, *p* < 0.001. For the proportion of movements on the easy set, we observed a significant main effect for movement, *F*(1,18) = 25.67, *p* < 0.001, no significant main effect of task, *F*(1,18) = 3.69, *p* = 0.071, and a significant interaction between the factors, *F*(1,18) = 6.08, *p* = 0.024. For the proportion of movements on the smaller set we observed significant main effects of movement, *F*(1,18) = 8.01, *p* = 0.011, as well as task, *F*(1,18) = 85.37, *p* < 0.001, and a significant interaction between the factors, *F*(1,18) = 43.92, *p* < 0.001. Finally, using the proportion movements on closer elements as dependent variable, we observed significant main effects of movement, *F*(1,18) = 227.78, *p* < 0.001, task, *F*(1,18) = 352.77, *p* < 0.001, and a significant interaction between factors, *F*(1,18) = 100.64, *p* < 0.001. For latencies, we observed significant main effects of the factors movement, *F*(1,18) = 14.68, *p* = 0.001, and task, *F*(1,18) = 167.81, *p* < 0.001, as well as a significant interaction between factors, *F*(1,18) = 6.21, *p* = 0.023.

In summary, participants showed tendencies to optimize their information sampling behavior in both, oculomotor and manual search tasks. However, while the sampling bias towards elements from the set of the eventually chosen target was consistently strong across different movements in trials of the manual search task, the sampling biases in the oculomotor search task emerged only with the second gaze shift. Instead, participants in the oculomotor search task preferentially targeted whichever element was closest to the current fixation location. While this behavioral pattern might suggest that in the oculomotor search task, participants treated lengthy saccades as costly, and thus tried to minimize them, the proximity bias disappeared from the second gaze shift onwards. Thus, we interpret the observed proximity bias of first gaze shifts not as a consequence of costly gaze shifts to distant elements, but as a general heuristic to select whatever stimulus was closest to fixation as gaze shift target (see also Discussion).

### Oculomotor search performance is more strongly constrained by noise

In both tasks, participants showed tendencies to act in accordance with an ideal observer, who aims to maximize its accumulated bonus payout: participants considered both the relative search costs and target difficulty for decisions about which target to search for, and they preferentially sampled elements from the set of the eventually chosen target while searching. However, participants, especially in the oculomotor search task, did not act perfectly rational: they occasionally chose targets with lower monetary gain (Figure 3A–B & Figure S1–S2), followed simple heuristics during information sampling (Figure 4C&F), and occasionally fixated elements from the non-chosen set while searching (Figure 4A&D).

**Figure 4.**
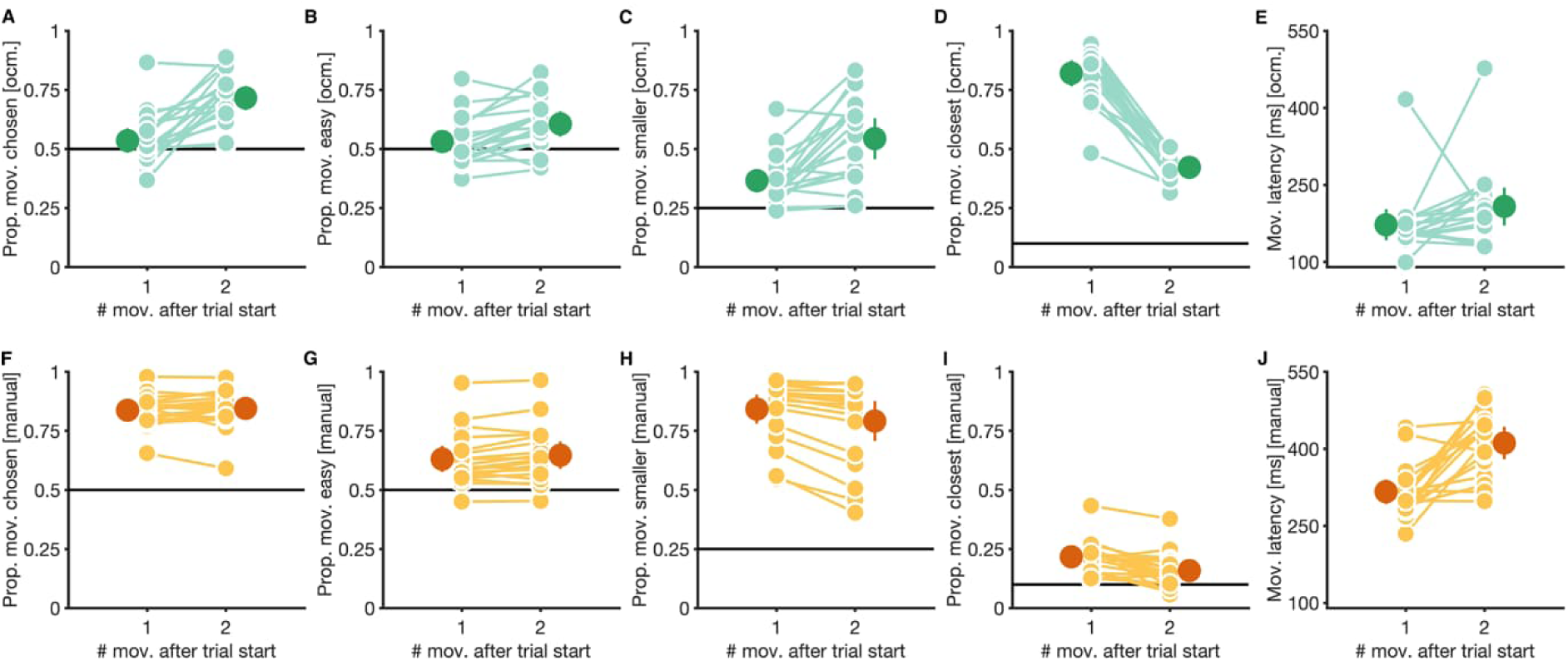
Behavior during information sampling. Proportion movements to elements from the set of the eventually chosen target (A&F), the easy set (B&G), the smaller set (C&H), and to elements closest to the current fixation (oculomotor search) or stylus location (manual search) (D&I) as well as latencies of first and second movements in trials (E&J). Data is shown separately for the oculomotor search (A–E) and manual search task (F–J). Small, light dots are data from individual participants, large, dark dots are means. Error bars are 95% confidence intervals. Black horizontal lines mark chance levels for the respective panel.

Consequently, in the double-target condition of the oculomotor search task, participants (*M* = 0.97 Cent/s, *CI*_95%_ [0.84 Cent/s, 1.10 Cent/s]) performed worse than an ideal observer (*M* = 1.45 Cent/s, *CI*_95%_ [1.28 Cent/s, 1.62 Cent/s]), who always choses higher-gain targets, and exclusively inspects elements from the set of the eventually chosen target during search (Figure 5A), *Y_t_*(12) = -9.28, *p* < 0.001. In contrast, in the manual search task, participants (*M* = 0.69 Cent/s, *CI*_95%_ [0.62 Cent/s, 0.75 Cent/s]) came close to the ideal observer’s performance (*M* = 0.75 Cent/s, *CI*_95%_ [0.70 Cent/s, 0.80 Cent/s]), *Y_t_*(12) = -2.54, *p* = 0.026 (Figure 5B).

**Figure 5.**
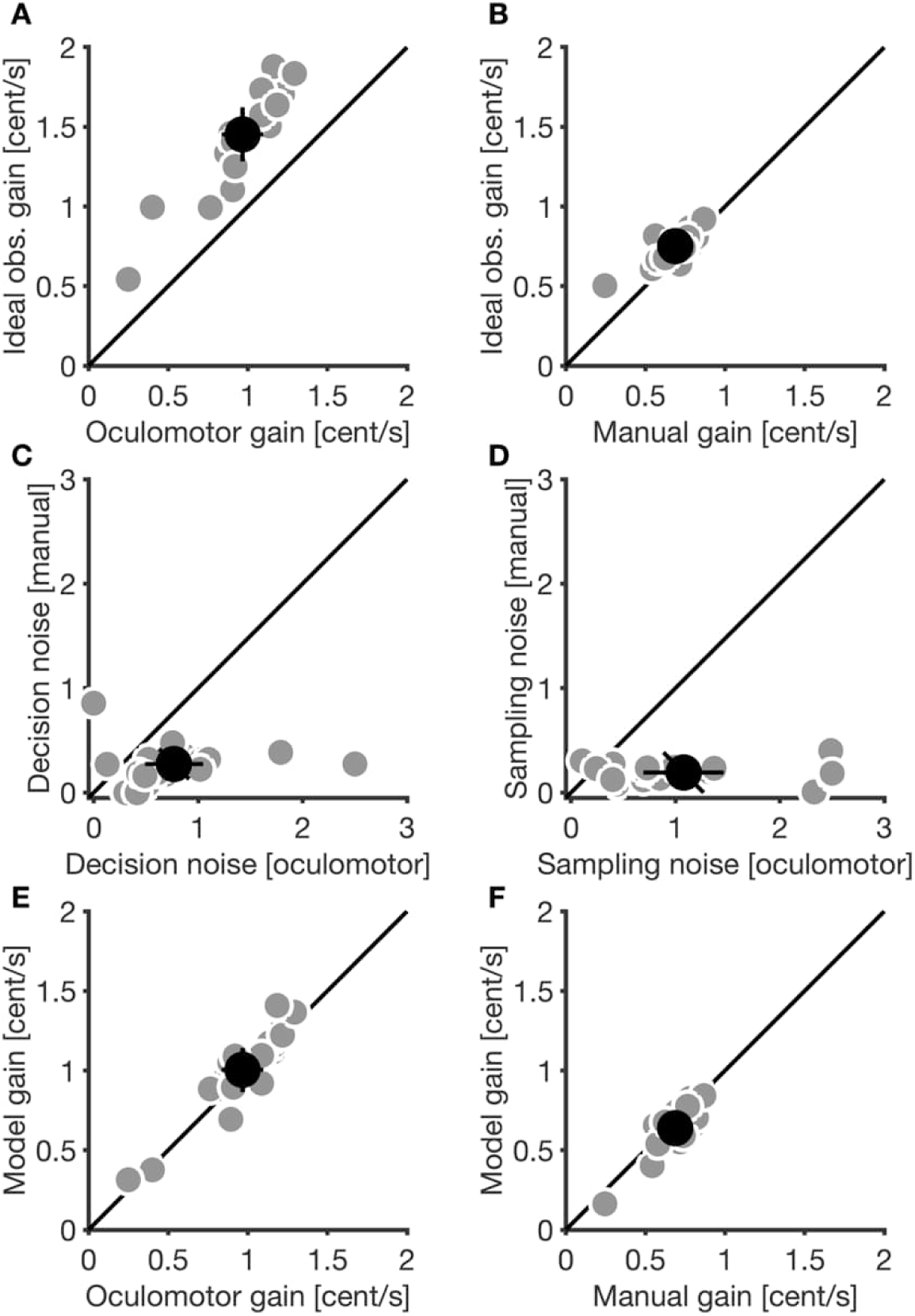
Model parameter as well as comparison between model predictions and empirical data in the double-target conditions of both tasks. Comparison between ideal observer gain and empirical gain (A&B), comparison between decision- and sampling-noise (C&D), and comparison between gain, as predicted by the generative stochastic model, and empirical gain. Small, light dots are data from individual participants, large, dark dots are means. Error bars are 95% confidence intervals.

One reason for the observed deviations from the behavior of an ideal observer might be that participant’s decision-making behavior was corrupted by noise at two distinct information processing levels: decision-noise might have corrupted choices between targets, while sampling-noise might have corrupted decisions for which elements to inspect during search. Due to the different time course of visual and manual actions, the magnitude of this decision and sampling-noise might have differed between tasks, leading to stronger deviations from the ideal observer performance in the oculomotor search compared to the manual search task. To test for this, we fitted a generative stochastic model (18) separately to the single-subject data of each task, and inspected the noise parameter of the model (see Methods for details).

In line with this rational, we did indeed observe that both, decision-noise (oculomotor search: *M* = 0.77, *CI*_95%_ [0.49, 1.05]; manual search: *M* = 0.27, *CI*_95%_ [0.18, 0.36]) and sampling-noise (oculomotor search: *M* = 1.08, *CI*_95%_ [0.69, 1.46]; manual search: *M* = 0.19, *CI*_95%_ [0.15, 0.23]) influenced behavior in both tasks (Figure 5C–D). However, decision-noise, *Y_t_*(12) = 5.53, *p* < 0.001, and sampling-noise, *Y_t_*(12) = 3.05, *p* = 0.010, contributed more strongly to behavior in the oculomotor search than the manual search task. Considering noise at different levels of decision making, our model was able to accurately predict participants empirical gain in the oculomotor search (*M* = 1.00 Cent/s, *CI*_95%_ [0.87 Cent/s, 1.14 Cent/s]), *r_s_* = 0.90, *p* < 0.001), and manual search task (*M* = 0.64 Cent/s, *CI*_95%_ [0.56 Cent/s, 0.71 Cent/s]), *r_s_* = 0.73, *p* = 0.001 (Figure 5E–F). Furthermore, our model also successfully captured decision-making behavior (Figure 6A–B, Figure S3–S4) as well information sampling behavior during search for targets (Figure 6C–D, Figure S5–S6), again, for both the oculomotor search (empirical proportion fixations on chosen elements: *M* = 0.70, *CI*_95%_ [0.66, 0.73]; predicted proportion choices on chosen elements: *M* = 0.69, *CI*_95%_ [0.67, 0.75]), *r_s_* = 0.97, *p* < 0.001, and manual search task (empirical proportion movements on chosen elements: *M* = 0.85, *CI*_95%_ [0.82, 0.88]; predicted proportion choices on chosen elements: *M* = 0.84, *CI*_95%_ [0.81, 0.87]), *r_s_* = 0.90, *p* < 0.001.

**Figure 6.**
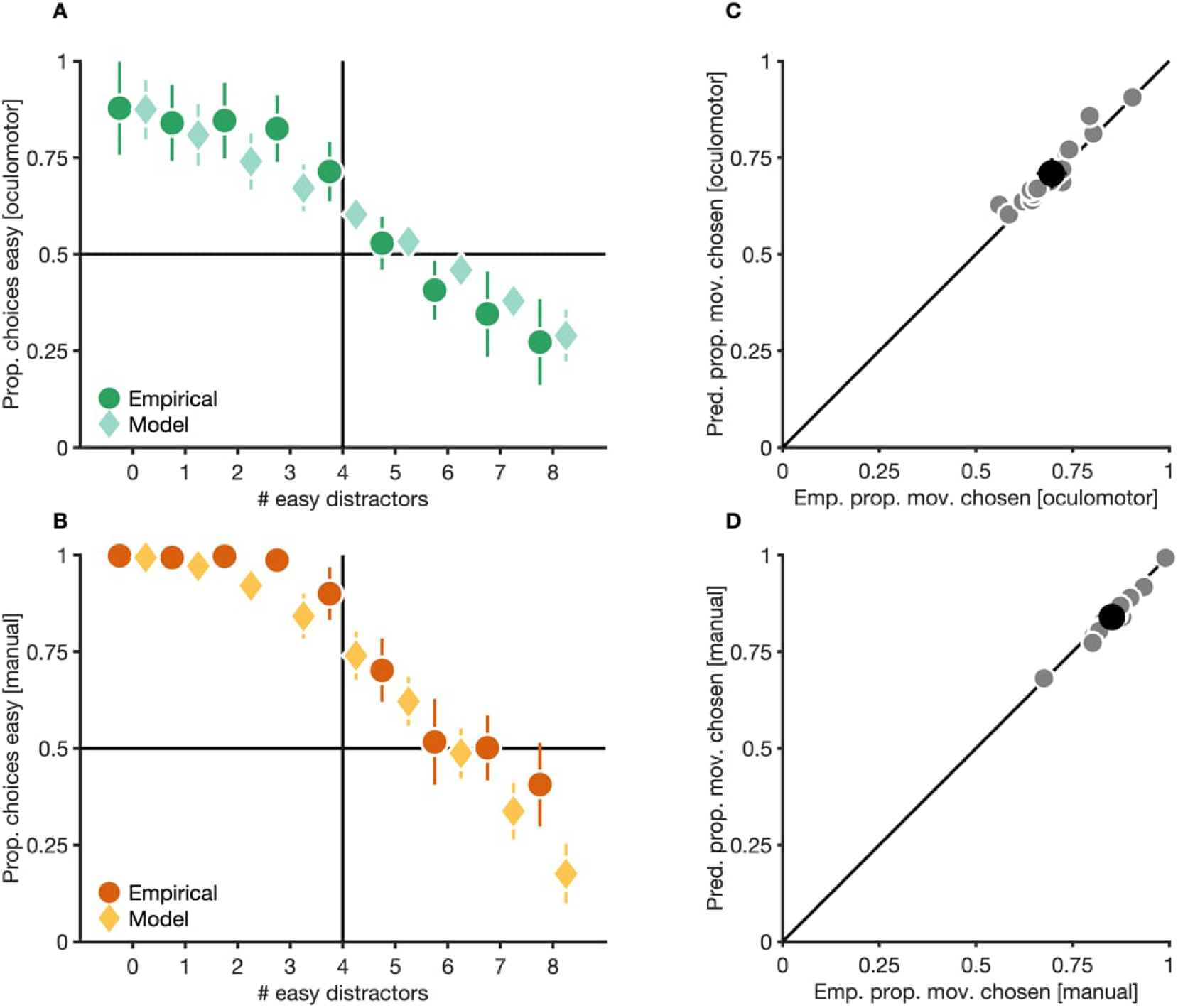
Comparison between empirical choice and information sampling behavior as well as choice and information sampling behavior, as predicted by the generative stochastic model. (A&B) Empirical and predicted probability to choose the easy target for different set sizes. To handle the proportional data, we transformed raw data in panels (A) & (B) with an arcsine transformation, calculated means and confidence intervals with the transformed values, and, finally, converted means and confidence intervals back to the original scale. (C–D) Empirical and predicted mean probability to target elements from the chosen set. Model predictions for the proportion of movements on the chosen set were calculated by considering the last movement immediately before a response. Since this last movement is not strictly part of the search process, it was omitted when calculating empirical proportion movements on the chosen set (see Figure 4A&D). To account for this difference, empirical proportion movements for this figure were calculate by taking the last movement in a trial (which mostly ended up on a target) into account. Single-subject proportions were calculated by averaging over the proportions for individual set-size conditions for each participant individually. Small, light dots are data from individual participants, large, dark dots are means. Error bars are 95% confidence intervals.

In summary, modeling revealed that both sampling and decision-noise influenced behavior in the two tasks. However, the contribution of both types of noise was greater in the oculomotor search task, explaining while participant’s behavior in this task showed greater deviations from the behavior of a theoretical ideal observer.

## Discussion

In two recent studies, we found evidence that humans trade off the search costs and perceptual benefits to optimize decision-making (18, 19). Here we studied whether this trade-off differs between two types of effectors: gaze shifts and manual actions. We instructed participants to either use gaze shifts (oculomotor search) or manual actions (stylus taps on a tablet; manual search) to search for targets that differed in their relative search costs and discrimination difficulty. Participants could freely choose between target options, and we incentivized them to optimize decision-making by limiting the total time they had to complete as many trials as possible and by rewarding participants based on their discrimination performance.

We found that participants, irrespective of which effector they used for search, showed some trade-off between search costs and discrimination difficulty for decision-making (Figure 3C). However, a comparison between the observed behavior of participants and the normative predictions of an ideal observer model revealed striking differences between effectors: While participants in the oculomotor search task occasionally chose lower-gain targets (Figure 3A) and tended to follow simple search heuristics, especially early in the search (Figure 4A&D), participants in the manual search task behaved closer to what was predicted by our ideal observer model (Figure 3B, Figure 4E&H). Consequently, while participants in the oculomotor search task underperformed compared to what the ideal observer model predicted (Figure 5A), participants in the manual search task came close to maximizing their accumulated reward over trials (Figure 5B). Using a generative stochastic model, we showed that this difference between effectors was likely due to a stronger contribution of decision-noise, corrupting decision about which target to search for, and sampling-noise, corrupting decisions about which information to sample during search, in the oculomotor search task (Figure 5C–F).

### Properties of eye movements make them more susceptible to noise

One reason for the observed differences between the oculomotor and manual search tasks might be related to some inherent properties of eye movements, which make them more susceptible to noise at different levels of decision making. Participants in our paradigm relied on saccadic eye movements to rapidly shift gaze between stimuli, while searching for targets. Saccades are ballistic movements because of their short duration (46), meaning that they cannot be corrected online. Furthermore, decisions between competing saccade targets are known to be modulated by the saccade latency, with (faster) lower-latency saccades being primarily driven by low-level stimulus features, while (slower) longer-latency saccades are under stronger top-down control (47–49). However, increasing the latency of saccades requires active inhibition of fast, reactive eye movements, which constitutes an effortful process (49).

Manual actions, such as reaching movements (the dominant type of movement in the manual search task), differ in various motor properties from saccades. For example, while saccades are short, fast and biomechanically cheap, reaching movements have greater temporal expenditure and require a greater expenditure of energy, resulting in a greater movement-inherent costs (29). Furthermore, while saccade trajectories cannot be corrected during ongoing movements, humans use online feedback about the quality of reaching movements to adjust the corresponding trajectories, even during ongoing movements. Interestingly, various studies found that humans consider the costs and benefits of ongoing manual actions, allowing for online updates of movement trajectories that reflect an evolving cost/benefit evaluation of competing reaching targets (50–54).

Thus, one reason for why behavior in the manual search task was less susceptible to noise might be related to the greater flexibility in controlling manual actions that unfold on a much slower time scale, and the comparatively large costs of inhibiting saccades for greater top-down oculomotor control. The greater temporal expenditure of manual actions, for example, might have given participants more time to evaluate search displays, and plan a course of action. Similarly, the ability to change the trajectories of manual actions online allows participants to dynamically update their chosen target based on a constantly progressing evaluation of the cost/benefit ratio of target options.

To avoid costly inhibition of eye movements, the oculomotor system, on the other hand, might have relied on simple behavioral heuristics early in search (9, 11, see also 30, 55), such as choosing the closest eye movement target. Under this interpretation, stimulus displays might have been evaluated during the ongoing visual search, and visual information sampling behavior became progressively more top-down driven (i.e., biased to elements from the eventually chosen set, from the second gaze shift onwards; Figure 4A). This transition from heuristics to top-down control would be in line with a progressing evaluation of the cost/benefit ratio of target options over a sequence of gaze shifts (56). However, unlike in the manual search task, search display evaluation in the oculomotor search task relied exclusively on visual information that was available after movement completion, without the benefit of being able to update movement trajectories online. This combination of a lower flexibility in movement execution and greater costs of movement inhibition might have resulted in sampling of more seemingly non-instrumental visual information.

One limitation of our study is that our setup only allowed us to measure endpoints of manual actions, but not their trajectories. Thus, our data does not allow for inferences about whether trajectories of manual actions were indeed updated online, whether manual actions were just easier to inhibit compared to gaze shifts, whether the greater temporal expenditure of manual actions allowed for more farsighted planning (29), or a combination of all those factors. However, while all those strategies would allow participants to reduce noise in manual decision-making, only inhibition of premature manual actions has the additional benefit of minimizing the number of costly manual actions.

### Eye movement behavior under resource constraints

One open question in the literature is whether eye movements are optimally tuned to the requirements of perceptual tasks (for reviews see 57, 58). While some studies have reported that human observers, in a variety of tasks, do indeed select saccade targets that maximize the gain in task-relevant visual information (59–66), other studies reported that human eye movement behavior is suboptimal (8, 16, 30–32, 67–74). Despite diverging results, all those studies have in common that they tested whether observers know about their own performance limitations as well as properties of their visual systems (e.g., its foveated nature), and exploit this knowledge to optimize target selection for eye movements. However, what most of those studies typically do not account for are the costs of eye movements, and how those costs might constrain behavior (for some notable exceptions, see 8, 9, 11, 16).

For example, Nowakowska et al. (70) asked participants to confirm the presence of a target in a stimulus grid, by sampling visual information via eye movements. In this task, the stimulus grid was split into two parts: one part was designed such that participants could confirm the target’s presence with peripheral vision, without the need to make any saccades to this part, while verifying the target’s presence in the other part of the grid required foveal vision and eye movements. Similar to our task, the optimal solution to the task of Nowakowska et al. (70) was to inhibit early saccades, evaluate the entire search grid via peripheral vision, and then direct eye movements to informative grid-locations only (i.e., one’s where the target’s presence could not be easily evaluated via peripheral vision). Despite the apparent simplicity of this task, the authors found that human observers consistently and repeatedly made eye movements to the uninformative grid-side, even though those eye movements did not help resolve uncertainty about the target’s presence and imposed substantial search costs by prolonging response times.

While inhibiting early saccades to suddenly appearing stimuli, and simultaneously using peripheral vision to plan a course of action, might be the optimal solution to a seemingly simple problem, this strategy is also computationally costly (for a review, see 75) and effortful (49). Partially relying on simple oculomotor heuristics, like choosing saccade targets closest to the current fixation location and/or saccade targets that satisfy inherent oculomotor preferences (9, 11), might be an attempt of the oculomotor system to produce goal-directed behavior with limited cognitive and computational resources (for reviews see 76–79). Fast reactions to suddenly appearing visual stimuli are also favored by the temporal discounting of reward (80, for reviews see 81, 74), which leads to lower reward when more time is required to foveate a target and which disincentivizes long deliberation before a saccade is executed.

Since saccades have short latencies and durations and are biomechanically cheap, heuristics-driven saccades can be quickly corrected by subsequent saccades to more informative locations, without imposing excessive additional temporal or energetic costs (but see 82). Furthermore, continuously updating eye movement planning over a series of fixation (56), while taking into account an evolving cost/benefit evaluation of targets, would allow for adaptive behavior in dynamic environments, without the need to commit to a fixed course of action after a computationally costly evaluation of visual input based on a single fixation. Overall, partially relying on heuristics to guide eye movements might yield a good enough performance, while simultaneously avoiding to impose computational costs that likely yield little behavioral benefit (see also, 16, 55, for a similar but more general argument see 83).

While heuristics-driven gaze shifts primarily occurred early in trials of the oculomotor search task, information sampling behavior later in trials was mostly characterized by instrumental gaze shifts to elements from the set of the eventually chosen target. However, even later in trials, some proportion of gaze shifts (and manual actions) sampled information that was non-instrumental for finding the chosen target. Is there a relevance to those seemingly superfluous movements later in trials? One possibility is that they are, in fact, not superfluous, but serve a functional purpose.

For example, Nowakowska et al. (70) suggested that participants might use eye movements to the uninformative grid-side to confirm and/or verify existing beliefs about the target’s presence. More generally, such a behavioral pattern might be part of an ongoing exploration-exploitation trade-off (84, 85), where a small proportion of actions is directed to random locations in the environment. Exploration-exploitation trade-offs are primarily relevant in complex environments (78), where they help experiencing novel states that an agent would otherwise not experience if it solely focused on maximizing (exploiting) some quantity. However, such trade-offs might also be helpful in the relatively simple laboratory environment of our task, where participants might randomly explore stimuli to verify existing beliefs about the relative discrimination difficulty of target options, despite sampling information from seemingly uninformative locations (for a review, see 86). While we did not directly quantify such trade-offs, the sampling-noise parameter in our generative stochastic model might implicitly capture such behavioral tendencies.

Overall, results from our study and other recent studies, which find evidence for an influence of oculomotor costs on eye movement behavior (9, 11, 16), highlight that the question is not necessarily whether eye movements are per se optimally tuned to task-requirements. The question is rather whether eye movement behavior yields a good-enough task-performance under computational constraints and in presence of inherent oculomotor costs.

### Interactions between gaze shifts and manual actions

The goal of our study was to compare decision-making behavior between gaze shifts and manual actions. However, since we did not simultaneously record data from both effectors, we cannot make any claims about the for interaction between gaze shifts and manual actions in the manual search task. For example, participants might have used gaze shifts to visually guide manual actions to whatever stimulus was chosen for inspection. This would be in line with results from studies on natural eye-hand-coordination, where eye movements were found to be made shortly before reaching movements towards task-relevant objects, and provided visual guidance to ensure accurate subsequent interactions (1, 87). In this case, gaze behavior in the manual search task might be completely determined by the choices of the manual movements and different from the gaze behavior in the oculomotor search task. Alternatively, gaze shifts and manual actions in the manual search task might, to some extent, be decoupled: For example, gaze shifts might initially be used to explore environments and plan a course of action, while manual actions are inhibited. Only after visual exploration is completed, manual actions are executed in accordance with the planned course of action, while gaze shifts are then used for visual guidance.

As in our paradigm, most studies that investigated differences and similarities between gaze shifts and manual actions in search paradigms either only measured data from both effectors in isolation (29, 55, 88) or used paradigms where gaze shifts and manual actions had no strong coupling beyond visual guidance (28, 89). For example, Moskowitz, Fooken, et al. (89) asked participants to visually search for targets and, after finding the target, to reach towards it’s on-screen location. The authors found that visual search was biased towards locations that would minimize the biomechanical costs of reaching towards the target’s locations after finding it. Thus, although it remains unclear how precisely eye movements and manual actions interact in multi-effector search tasks, the study by Moskowitz, Fooken, et al. (89) at least suggest that eye movements during visual search are not exclusively for visual guidance, but can also serve the later affordances of manual actions (see also 90).

Overall, given existing literature, it is plausible to assume that gaze shifts in the manual search task might have served a purpose beyond pure visual guidance and contributed to planning of manual actions. While visually exploring environments in absence of any manual actions might not yield any information about which stimuli are targets and which are distractors, such a behavior might, for example, be instrumental estimating the relative search costs of target options by sampling color information from individual stimuli.

## Conclusion

We found that human participants, irrespective of whether they use gaze shifts or manual actions to search for targets, traded off search costs and discrimination benefits of competing target options to optimize task-performance. However, movement inherent properties influenced the effector-specific manifestation of this trade-off, resulting not only in global behavioral similarities across effectors, but also in specific behavioral patterns that presumably echo the respective biomechanical and computational constraints of effectors.

## Declarations

### Funding

ACS, AS and IW were funded by the Deutsche Forschungsgemeinschaft (DFG, German Research Foundation) – project number 222641018 – SFB/TRR 135 TP B2 and TP B3 and the DFG International Research Training Group IRTG-1901-The Brain in Action. ACS, AS, and JT received funding from the Excellence Program of the Hessian Ministry of Higher Education, Science, Research and Art (“The Adaptive Mind”). IW received additional funding from by Meta Platforms, Inc.

### Conflicts of interest

The authors have no relevant financial or non-financial interests to disclose.

### Ethics approval

The experiment was conducted in accordance with the ethical guidelines laid down in the 1964 declaration of Helsinki and was approved by the ethics committee of the Marburg University, Department of Psychology (proposal 2017-27k).

### Consent to participate

All participants provided informed consent prior to testing.

### Consent for publication

All participants provided informed consent that their data will be published in a scientific article.

### Authors’ contribution

Experimental design: ACS, AS, IW, JT; data collection: IW, JT; data analysis: IW (preprocessing & modelling), JT (preprocessing); data visualization: IW; writing first draft: IW; revising manuscript: IW, ACS, AS, JT.

## Acknowledgements

We thank Joni Blume and Christoph Schäfer for helping with data collection.

## Supplementary Information

### Supplementary figures

**Supplementary figure S 1.**
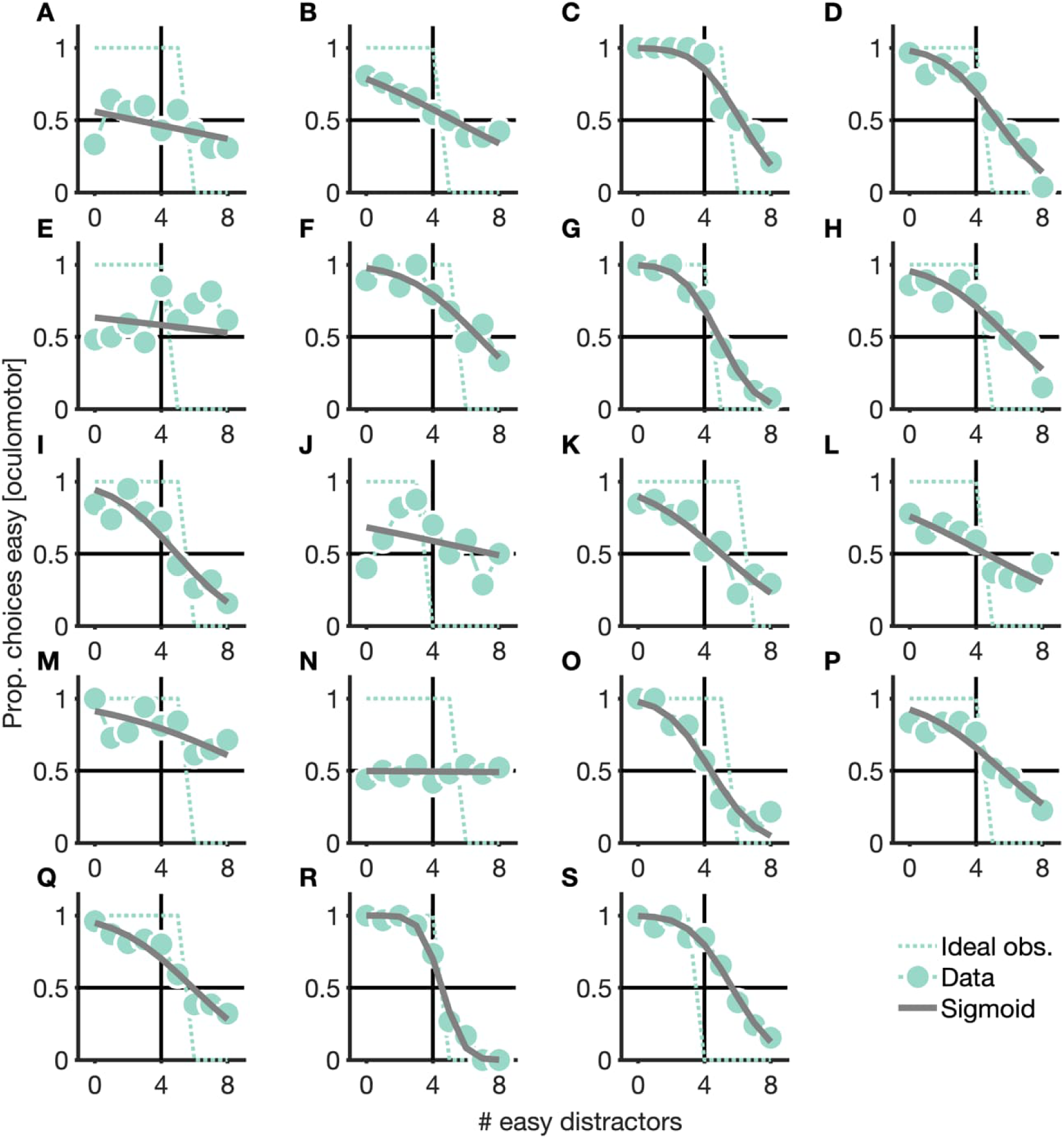
Empirical and ideal observer choice behavior of participants in the oculomotor search task. (A–S) Proportion of trials in which individual participants in the double-target condition of the oculomotor search task chose to discriminate easy targets. Small, light dots are data, gray solid lines are fitted linear regressions, light dotted lines are ideal observer predictions.

**Supplementary figure S 2.**
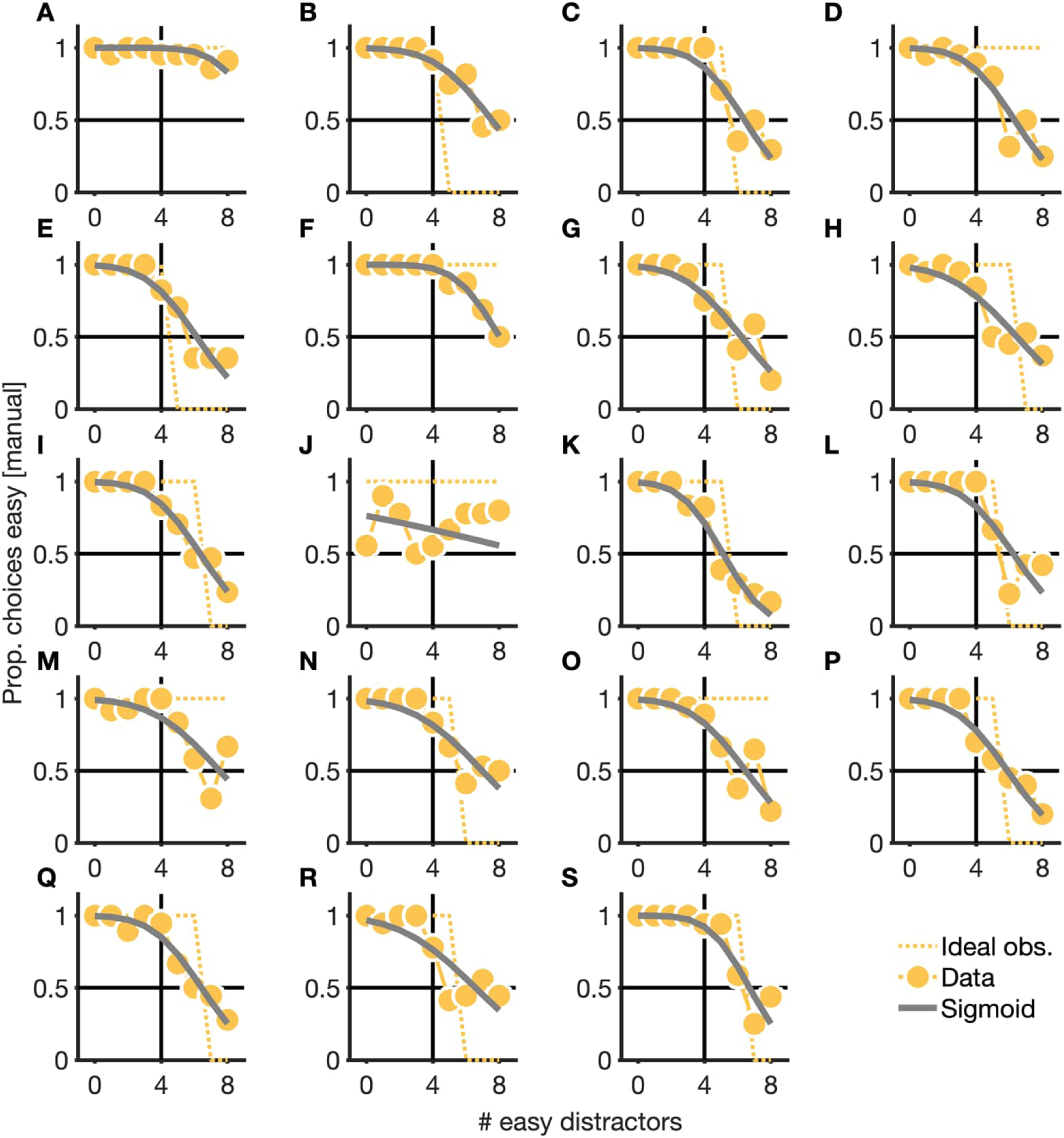
Empirical and ideal observer choice behavior of participants in the manual search task. (A–S) Proportion of trials in which individual participants in the double-target condition of the manual search task chose to discriminate easy targets. Small, light dots are data, gray solid lines are fitted linear regressions, light dotted lines are ideal observer predictions.

**Supplementary figure S 3.**
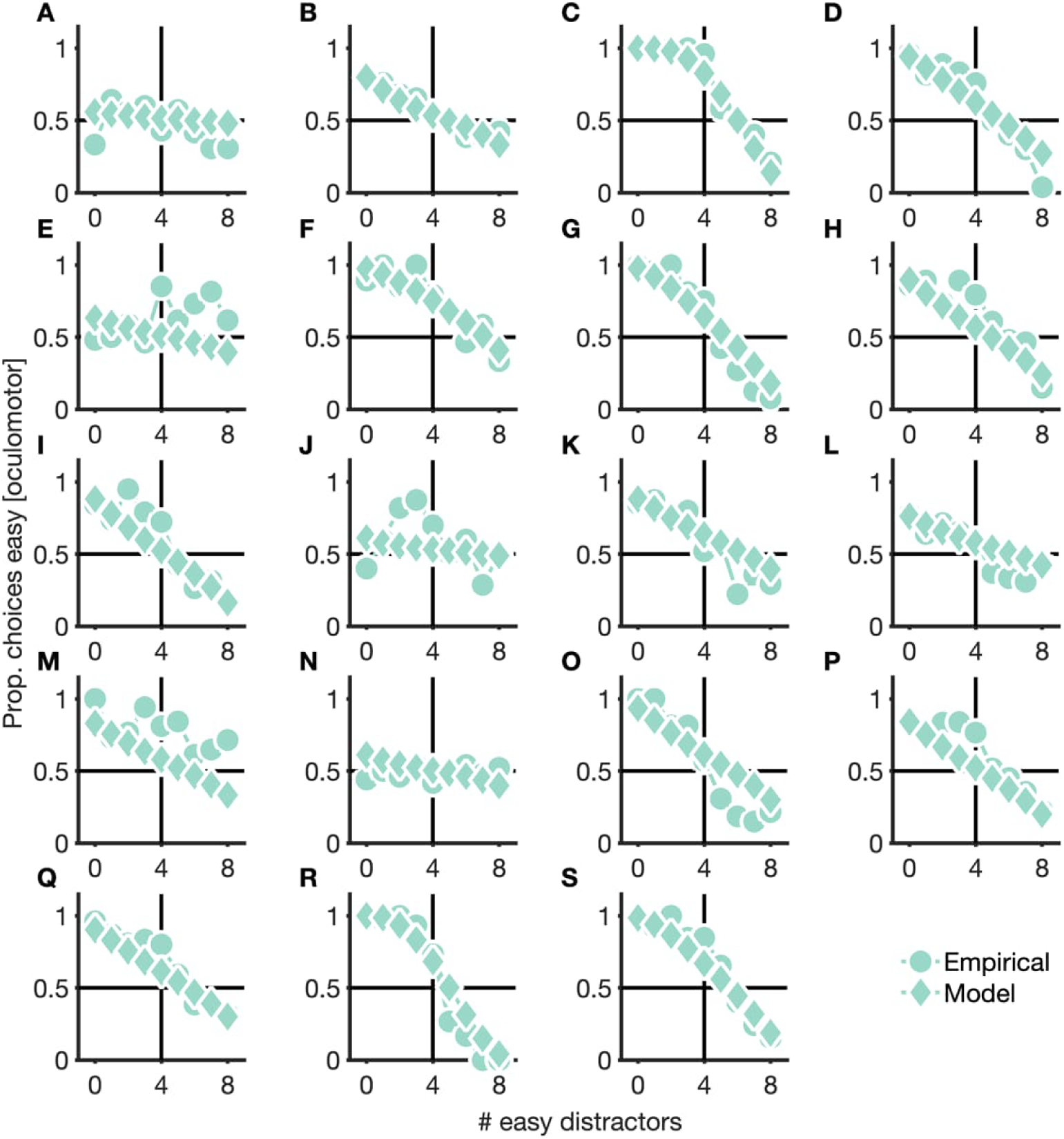
Empirical choice behavior and choice behavior, as predicted by the generative stochastic model, of participants in the oculomotor search task. (A–S) Proportion of trials in which individual participants in the double-target condition of the manual search task chose to discriminate easy targets. Light dots are data, light diamonds are model predictions.

**Supplementary figure S 4.**
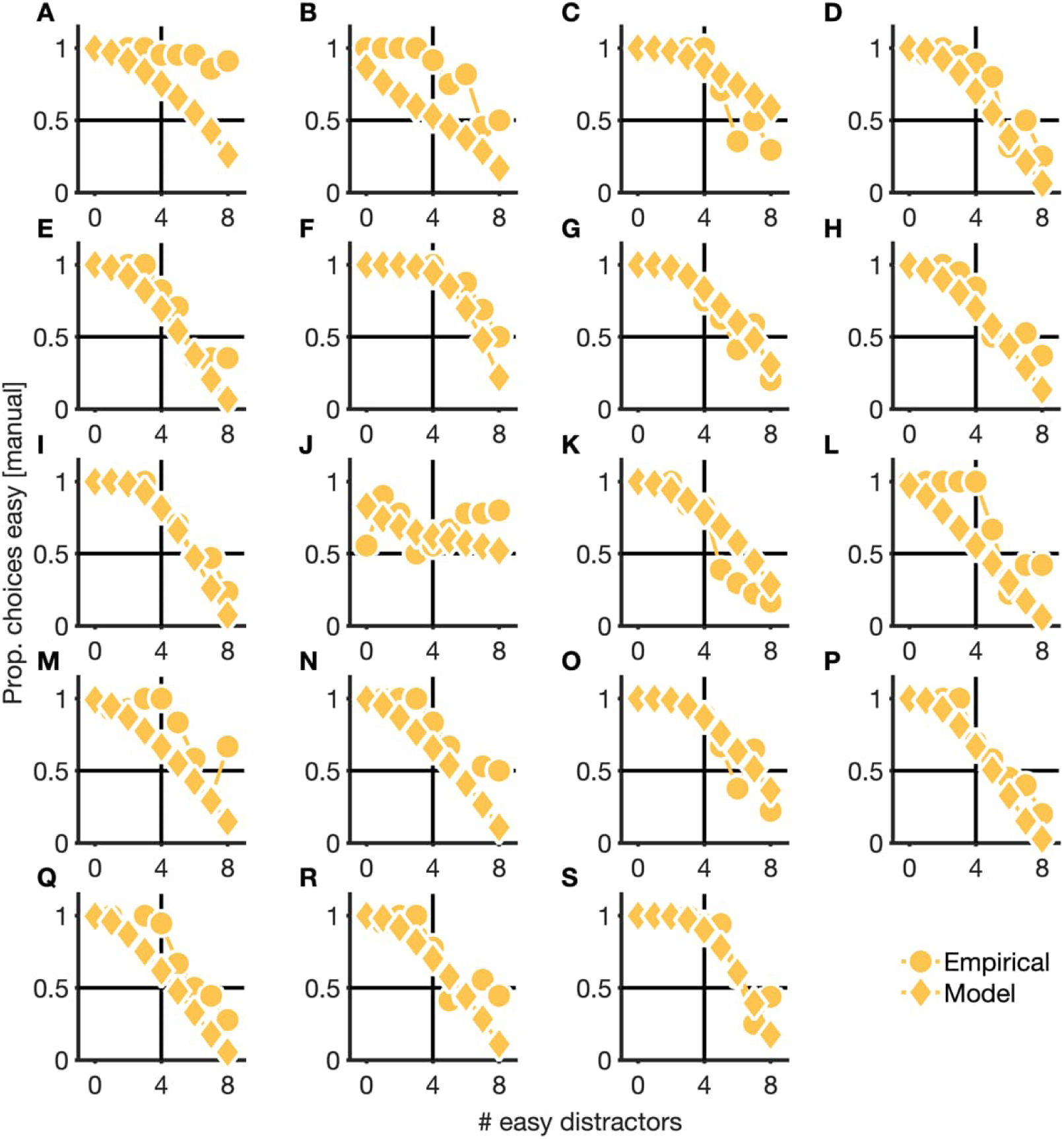
Empirical choice behavior and choice behavior, as predicted by our generative stochastic model, of participants in the manual search task. (A– S) Proportion of trials in which individual participants in the double-target condition of the manual search task chose to discriminate easy targets. Light dots are data, light diamonds are model predictions.

**Supplementary figure S 5.**
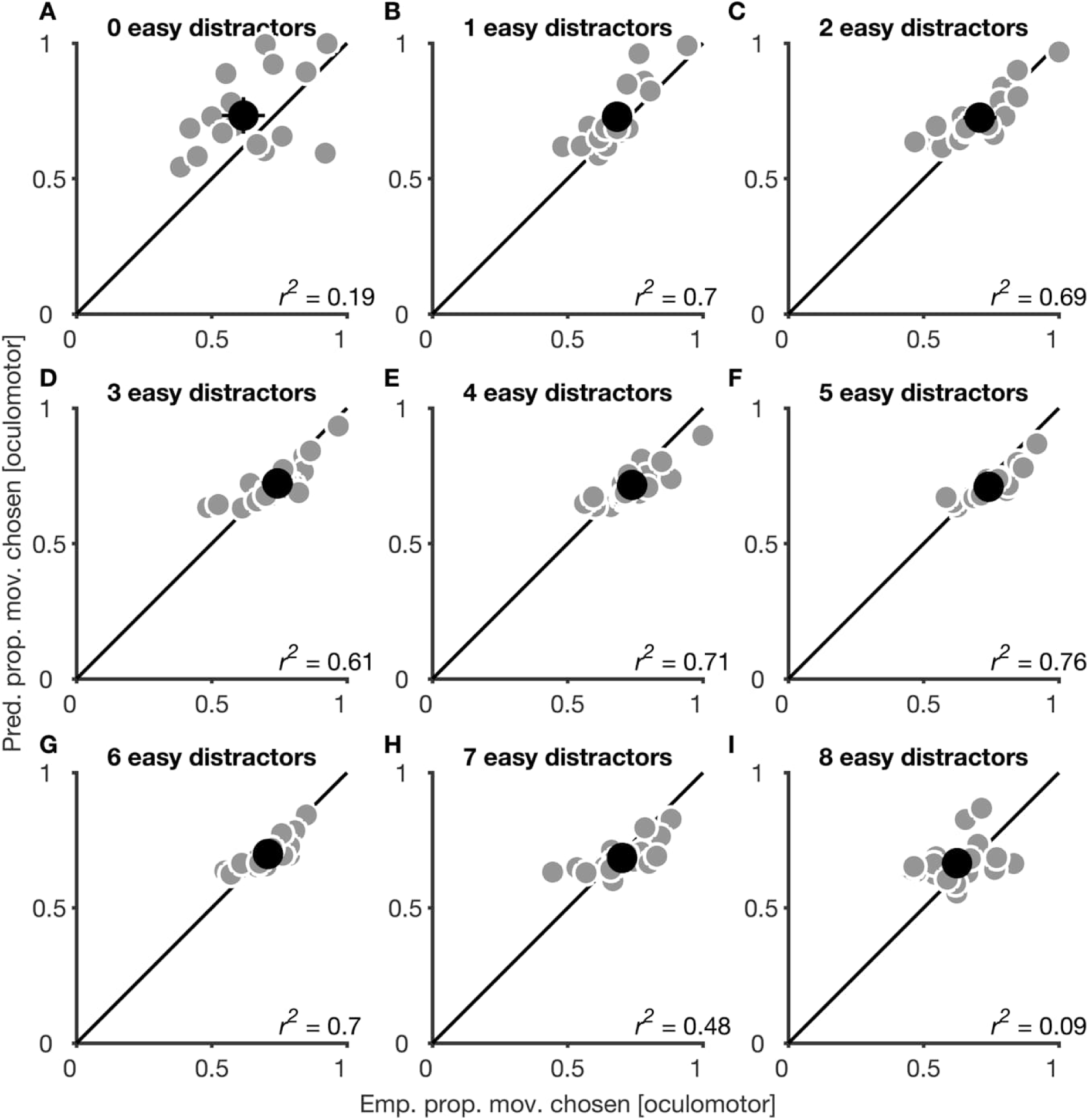
Empirical and predicted proportion of movements to elements from the chosen set, separately for each set-size condition in the oculomotor search task. See Figure 6 in the main manuscript for information about how proportion movements were calculated. Small, light dots are data from individual participants, large, dark dots are means. Error bars are 95% confidence intervals.

**Supplementary figure S 6.**
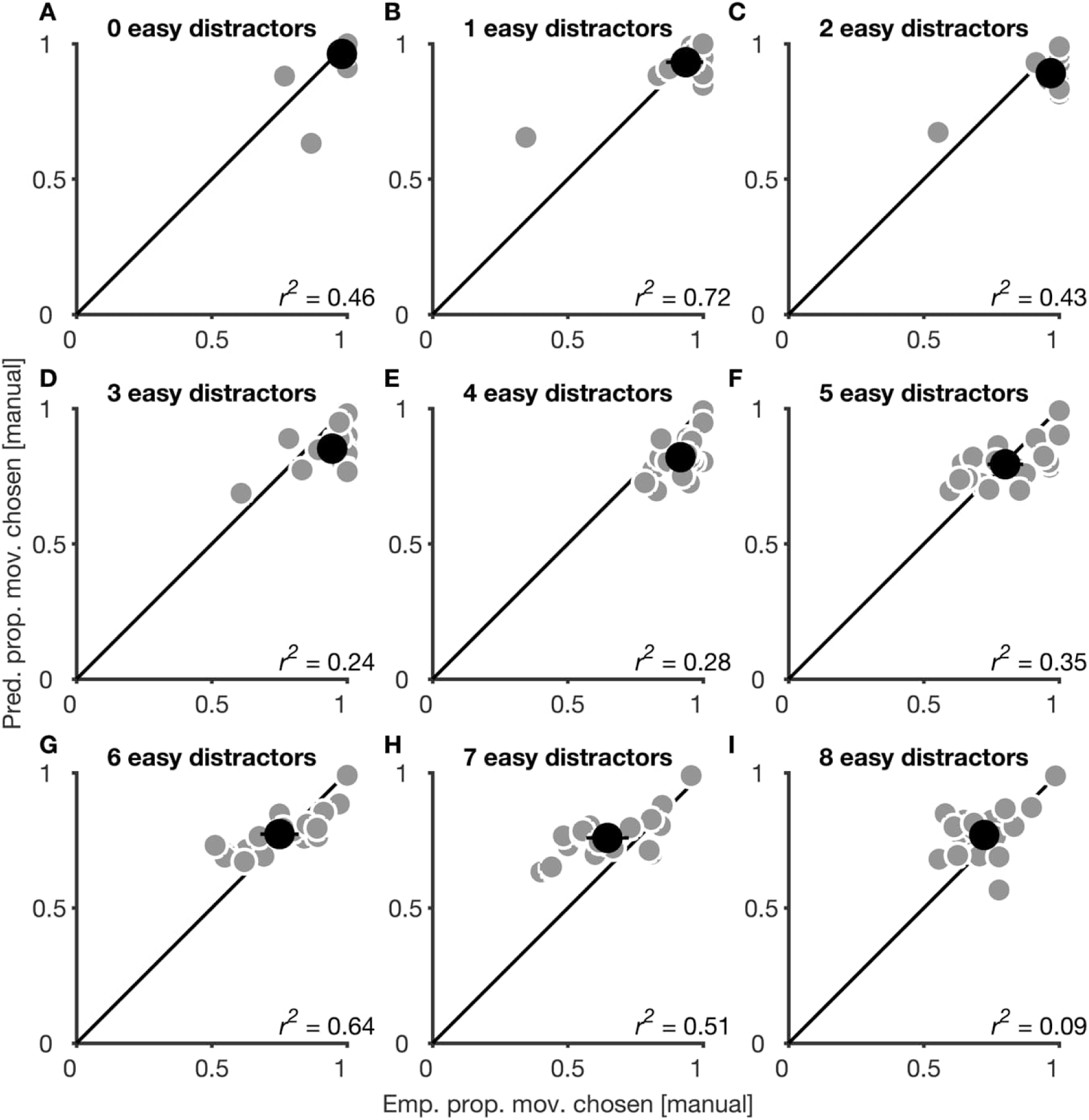
Empirical and predicted proportion of movements to elements from the chosen set, separately for each set-size condition in the manual search task. See Figure 6 in the main manuscript for information about how proportion movements were calculated. Small, light dots are data from individual participants, large, dark dots are means. Error bars are 95% confidence intervals.

## Appendix X: Trial exclusion due to “pen dragging”

*Participants were instructed to lift the pen from the tablet surface when moving from one stimulus to the next. However, occasionally participants dragged the pen across the surface, which could uncover stimuli but stylus down and up events were not reliably logged. This is problematic, because stylus down and up events were used to determine movement on- and offsets. We checked the trials against three criteria that indicate pen dragging:*

### C1: Stylus never released from fixation cross

*This criterion is fulfilled if no stylus up event was recorded for the initial pressing on the fixation cross. This happens when the stylus was dragged away from it and lifted elsewhere*.

### C2: In consistent order of stylus up / down events

*This criterion is fulfilled when the order of stylus down and up events for a stimulus was inconsistent. In particular, we checked whether the first encounter of the stimulus was a stylus down event, whether the number of stylus down and up events matched, and whether down and up alternated. If any of these were not true, the sequence is considered inconsistent, which can only happen if pen dragging uncovered stimuli. There was one exception: the stimulus encountered right before the response did not have to fulfill these criteria, because it was allowed to enter the response without lifting the pen first (a case in which violation of the criteria above are expected)*.

### C3: Target never uncovered

*This criterion was fulfilled when the response was entered without a recording of a stylus down event for one of the targets. In this case it is likely that the participant viewed the target stimulus via pen dragging*.

*A trial was flagged for removal if at least one of these criteria was fulfilled. It cannot be entirely excluded that there were some trials with undetected pen that did not produce conspicuous events. However, if it occurred, it could only have happened rarely as systematic use of pen dragging would have been detected with the criteria above. The table below lists the number of detected pen dragging trials and the criteria they fulfilled for every participant:*

| Participant | C1 | C2 | C3 | Trials Excl. | % Excluded |
| --- | --- | --- | --- | --- | --- |
| 1 | 0 | 2 | 1 | 3 | 0.83% |
| 2 | 0 | 16 | 0 | 16 | 8.25% |
| 4 | 0 | 1 | 1 | 2 | 0.65% |
| 6 | 0 | 16 | 13 | 29 | 11.60% |
| 7 | 5 | 4 | 4 | 10 | 3.76% |
| 8 | 0 | 1 | 0 | 1 | 0.34% |
| 9 | 0 | 1 | 1 | 2 | 0.73% |
| 13 | 0 | 4 | 0 | 4 | 1.31% |
| 14 | 1 | 24 | 5 | 29 | 12.66% |
| 16 | 1 | 6 | 3 | 9 | 3.41% |
| 17 | 0 | 2 | 0 | 2 | 0.65% |
| 19 | 98 | 0 | 259 | 259 | 77.78% |
| 20 | 0 | 6 | 0 | 6 | 2.01% |
| 21 | 0 | 0 | 2 | 2 | 0.76% |
| 24 | 0 | 7 | 0 | 7 | 2.66% |

## Notes

### Competing Interest Statement

The authors have declared no competing interest.

## References

1. Land M, Mennie N, Rusted J. The roles of vision and eye movements in the control of activities of daily living. Perception 28: 1311–1328, 1999. doi: 10.1068/p2935.

2. Schütz AC, Braun DI, Gegenfurtner KR. Eye movements and perception: A selective review. Journal of Vision 11: 1–30, 2011. doi: 10.1167/11.5.1.

3. Theeuwes J. Automatic control of visual selection. Nebr Symp Motiv 59: 23–62, 2012. doi: 10.1007/978-1-4614-4794-8_3.

4. Dodge R. Visual perception during eye movement. Psychological Review 7: 454–465, 1900. doi: 10.1037/h0067215.

5. Schweitzer R, Rolfs M. Intrasaccadic motion streaks jump-start gaze correction. Science Advances 7: 15, 2021. doi: 10.1126/sciadv.abf2218.

6. Hoppe D, Rothkopf CA. Learning rational temporal eye movement strategies. Proceedings of the National Academy of Sciences 113: 8332–8337, 2016. doi: 10.1073/pnas.1601305113.

7. Abekawa N, Gomi H, Diedrichsen J. Gaze control during reaching is flexibly modulated to optimize task outcome. Journal of Neurophysiology 126: 816–826, 2021. doi: 10.1152/jn.00134.2021.

8. Araujo C, Kowler E, Pavel M. Eye movements during visual search: The costs of choosing the optimal path. Vision Research 41: 3613–3625, 2001. doi: 10.1016/S0042-6989(01)00196-1.

9. Kadner F, Thomas T, Hoppe D, Rothkopf CA. Improving saliency models’ predictions of the next fixation with humans’ intrinsic cost of gaze shifts. In: 2023 IEEE/CVF Winter Conference on Applications of Computer Vision (WACV). 2023 IEEE/CVF Winter Conference on Applications of Computer Vision (WACV). IEEE, p. 2103–2113.

10. Lisi M, Solomon JA, Morgan MJ. Gain control of saccadic eye movements is probabilistic. Proceedings of the National Academy of Sciences 116: 16137–16142, 2019. doi: 10.1073/pnas.1901963116.

11. Thomas T, Hoppe D, Rothkopf CA. The neuroeconomics of individual differences in saccadic decisions. Neuroscience.

12. Ballard DH, Hayhoe MM, Pelz JB. Memory representations in natural tasks. Journal of Cognitive Neuroscience 7: 66–80, 1995. doi: 10.1162/jocn.1995.7.1.66.

13. Melnik A, Schüler F, Rothkopf CA, König P. The World as an External Memory: The Price of Saccades in a Sensorimotor Task. Front Behav Neurosci 12: 253, 2018. doi: 10.3389/fnbeh.2018.00253.

14. Schut MJ, Van der Stoep N, Postma A, Van der Stigchel S. The cost of making an eye movement: A direct link between visual working memory and saccade execution. Journal of Vision 17: 15, 2017. doi: 10.1167/17.6.15.

15. Somai RS, Schut MJ, Van der Stigchel S. Evidence for the world as an external memory: A trade-off between internal and external visual memory storage. Cortex 122: 108–114, 2020. doi: 10.1016/j.cortex.2018.12.017.

16. Zhou Y, Yu Y. Human visual search follows a suboptimal Bayesian strategy revealed by a spatiotemporal computational model and experiment. Communications Biology 4: 1–16, 2021. doi: 10.1038/s42003-020-01485-0.

17. Irons JL, Leber AB. Choosing attentional control settings in a dynamically changing environment. *Attention*, Perception, and Psychophysics 78: 2031–2048, 2016. doi: 10.3758/s13414-016-1125-4.

18. Wagner I, Henare D, Tünnermann J, Schubö A, Schütz AC. Humans trade off search costs and accuracy in a combined visual search and perceptual task. .

19. Henare DT, Tünnermann J, Wagner I, Schütz AC, Schubö A. Complex trade-offs in a dual-target visual search task are indexed by lateralised ERP components. Sci Rep 14: 22839, 2024. doi: 10.1038/s41598-024-72811-3.

20. Lio G, Fadda R, Doneddu G, Duhamel JR, Sirigu A. Digit-tracking as a new tactile interface for visual perception analysis. Nature Communications 10, 2019. doi: 10.1038/s41467-019-13285-0.

21. Yang Y, Mo L, Lio G, Huang Y, Perret T, Sirigu A, Duhamel J-R. Assessing the allocation of attention during visual search using digit-tracking, a calibration-free alternative to eye tracking. Sci Rep 13: 2376, 2023. doi: 10.1038/s41598-023-29133-7.

22. Brenner E, Smeets JBJ. Quickly making the correct choice. Vision Research 113: 198–210, 2015. doi: 10.1016/j.visres.2015.03.028.

23. Cos I, Bélanger N, Cisek P. The influence of predicted arm biomechanics on decision making. Journal of Neurophysiology 105: 3022–3033, 2011. doi: 10.1152/jn.00975.2010.

24. Cos I, Medleg F, Cisek P. The modulatory influence of end-point controllability on decisions between actions. Journal of Neurophysiology 108: 1764–1780, 2012. doi: 10.1152/jn.00081.2012.

25. Cos I, Duque J, Cisek P. Rapid prediction of biomechanical costs during action decisions. Journal of Neurophysiology 112: 1256–1266, 2014. doi: 10.1152/jn.00147.2014.

26. Hagura N, Haggard P, Diedrichsen J. Perceptual decisions are biased by the cost to act. eLife 6: e18422, 2017. doi: 10.7554/eLife.18422.

27. Marcos E, Cos I, Girard B, Verschure PFMJ. Motor Cost Influences Perceptual Decisions. PLoS ONE 10: e0144841, 2015. doi: 10.1371/journal.pone.0144841.

28. Moskowitz JB, Berger SA, Fooken J, Castelhano MS, Gallivan JP, Flanagan JR. The influence of movement-related costs when searching to act and acting to search. Journal of Neurophysiology 129: 115–130, 2023. doi: 10.1152/jn.00305.2022.

29. Diamond JS, Wolpert DM, Flanagan JR. Rapid target foraging with reach or gaze: The hand looks further ahead than the eye. PLOS Computational Biology 13: 1–23, 2017. doi: 10.1371/journal.pcbi.1005504.

30. Clarke ADF, Green P, Chantler MJ, Hunt AR. Human search for a target on a textured background is consistent with a stochastic model. Journal of Vision 16: 1–16, 2016. doi: 10.1167/16.7.4.

31. Clarke ADF, Nowakowska A, Hunt AR. Visual search habits and the spatial structure of scenes. Atten Percept Psychophys 84: 1874–1885, 2022. doi: 10.3758/s13414-022-02506-2.

32. Nowakowska A, Clarke ADF, von Seth J, Hunt AR. Search strategies improve with practice, but not with time pressure or financial incentives. Journal of experimental psychology Human perception and performance 47: 1009–1021, 2021. doi: 10.1037/xhp0000912.

33. The MathWorks Inc. The MathWorks Inc.: 2016. https://www.mathworks.com.

34. Brainard DH. The Psychophysics Toolbox. Spatial Vision 10: 433–436, 1997. doi: 10.1163/156856897X00357.

35. Kleiner M, Brainard D, Pelli D. What’s new in Psychtoolbox-3? [Online]. 2007. https://github.com/neurodebian/psychtoolbox-3-old-gitsvn-based/blob/master/Psychtoolbox/PsychDocumentation/Psychtoolbox3-Slides.pdf [23 Feb. 2023].

36. Pelli DG. The VideoToolbox software for visual psychophysics: transforming numbers into movies. Spat Vis 10: 437–442, 1997. doi: 10.1163/156856897X00366.

37. Cornelissen FW, Peters EM, Palmer J. The Eyelink toolbox: Eye tracking with MATLAB and the Psychophysics Toolbox. Behavior Research Methods, Instruments, & Computers 34: 613–617, 2002. doi: 10.3758/BF03195489.

38. Mathôt S, Schreij D, Theeuwes J. OpenSesame: An open-source, graphical experiment builder for the social sciences. Behav Res 44: 314–324, 2012. doi: 10.3758/s13428-011-0168-7.

39. Peirce JW. PsychoPy—Psychophysics software in Python. Journal of Neuroscience Methods 162: 8–13, 2007. doi: 10.1016/j.jneumeth.2006.11.017.

40. Thaler L, Schütz AC, Goodale MA, Gegenfurtner KR. What is the best fixation target? The effect of target shape on stability of fixational eye movements. Vision Research 76: 31–42, 2013. doi: 10.1016/j.visres.2012.10.012.

41. Irons JL, Leber AB. Characterizing individual variation in the strategic use of attentional control. Journal of Experimental Psychology: Human Perception and Performance 44: 1637–1654, 2018. doi: 10.1037/xhp0000560.

42. Yuen KK. The two-sample trimmed t for unequal population variances. Biometrika 61: 165e170, 1974. doi: 10.1093/biomet/61.1.165.

43. Wilcox RR. Introduction to robust estimation and hypothesis testing. 5th edition. Waltham, MA: Elsevier, 2022.

44. R Core Team. R: A language and environment for statistical computing [Online]. R Foundation for Statistical Computing: 2021. https://www.R-project.org/.

45. JASP Team. JASP. 2024.

46. Bahill AT, Clark MR, Stark L. The main sequence, a tool for studying human eye movements. Mathematical Biosciences 24: 191–204, 1975. doi: 10.1016/0025-5564(75)90075-9.

47. Schütz AC, Trommershäuser J, Gegenfurtner KR. Dynamic integration of information about salience and value for saccadic eye movements. Proceedings of the National Academy of Sciences 109: 7547–7552, 2012. doi: 10.1073/pnas.1115638109.

48. van Heusden E, van Zoest W, Donk M, Olivers CNL. An attentional limbo: Saccades become momentarily non-selective in between saliency-driven and relevance-driven selection. .

49. Wolf C, Lappe M. Top-down control of saccades requires inhibition of suddenly appearing stimuli. *Attention*, Perception, and Psychophysics 82: 3863–3877, 2020. doi: 10.3758/s13414-020-02101-3.

50. Cos I, Pezzulo G, Cisek P. Changes of Mind after Movement Onset Depend on the State of the Motor System. eNeuro 8: ENEURO.0174-21.2021, 2021. doi: 10.1523/ENEURO.0174-21.2021.

51. Lepora NF, Pezzulo G. Embodied Choice: How Action Influences Perceptual Decision Making. PLoS Comput Biol 11: e1004110, 2015. doi: 10.1371/journal.pcbi.1004110.

52. Michalski J, Green AM, Cisek P. Reaching decisions during ongoing movements. Journal of Neurophysiology 123: 1090–1102, 2020. doi: 10.1152/jn.00613.2019.

53. Nashed JY, Crevecoeur F, Scott SH. Rapid Online Selection between Multiple Motor Plans. J Neurosci 34: 1769–1780, 2014. doi: 10.1523/JNEUROSCI.3063-13.2014.

54. Pierrieau E, Lepage J-F, Bernier P-M. Action Costs Rapidly and Automatically Interfere with Reward-Based Decision-Making in a Reaching Task. eNeuro 8: ENEURO.0247-21.2021, 2021. doi: 10.1523/ENEURO.0247-21.2021.

55. Rajsic J, Wilson DE, Pratt J. The price of information: Increased inspection costs reduce the confirmation bias in visual search. Quarterly Journal of Experimental Psychology 71: 832–849, 2018. doi: 10.1080/17470218.2016.1278249.

56. De Vries JP, Hooge ITC, Verstraten FAJ. Saccades Toward the Target Are Planned as Sequences Rather Than as Single Steps. Psychological Science 25: 215–223, 2014. doi: 10.1177/0956797613497020.

57. Braun DI, Schütz AC. Eye Movements and Perception. Oxford Research Encyclopedia of Psychology.: 2022.

58. Wolf C, Lappe M. Vision as oculomotor reward: cognitive contributions to the dynamic control of saccadic eye movements. Cognitive Neurodynamics 15: 547–568, 2021. doi: 10.1007/s11571-020-09661-y.

59. Eckstein MP, Schoonveld W, Zhang S, Mack SC, Akbas E. Optimal and human eye movements to clustered low value cues to increase decision rewards during search. Vision Research 113: 137–154, 2015. doi: 10.1016/j.visres.2015.05.016.

60. Hoppe D, Rothkopf CA. Multi-step planning of eye movements in visual search. Scientific Reports 9: 1–12, 2019. doi: 10.1038/s41598-018-37536-0.

61. Najemnik J, Geisler WS. Optimal eye movement strategies in visual search. Nature 434: 387–391, 2005. doi: 10.1038/nature03390.

62. Najemnik J, Geisler WS. Eye movement statistics in humans are consistent with an optimal search strategy. Journal of Vision 8: 1–14, 2008. doi: 10.1167/8.3.4.

63. Paulun VC, Schütz AC, Michel MM, Geisler WS, Gegenfurtner KR. Visual search under scotopic lighting conditions. Vision Research 113: 155–168, 2015. doi: 10.1016/j.visres.2015.05.004.

64. Peterson MF, Eckstein MP. Looking just below the eyes is optimal across face recognition tasks. Proceedings of the National Academy of Sciences 109: E3314–E3323, 2012. doi: 10.1073/pnas.1214269109.

65. Renninger LW, Verghese P, Coughlan J. Where to look next? Eye movements reduce local uncertainty. Journal of Vision 7: 6, 2007. doi: 10.1167/7.3.6.

66. Yang SC-H, Lengyel M, Wolpert DM. Active sensing in the categorization of visual patterns. eLife 5: e12215, 2016. doi: 10.7554/eLife.12215.

67. Ackermann JF, Landy MS. Choice of saccade endpoint under risk. Journal of Vision 13: 27–27, 2013. doi: 10.1167/13.3.27.

68. Ghahghaei S, Verghese P. Efficient saccade planning requires time and clear choices. Vision Research 113: 125–136, 2015. doi: 10.1016/j.visres.2015.05.006.

69. Morvan C, Maloney LT. Human visual search does not maximize the post-saccadic probability of identifying targets. PLOS Computational Biology 8: e1002342–11, 2012.

70. Nowakowska A, Clarke ADF, Hunt AR. Human visual search behaviour is far from ideal. Proceedings of the Royal Society B: Biological Sciences 284: 20162767, 2017. doi: 10.1098/rspb.2016.2767.

71. Tsank Y, Eckstein MP. Domain Specificity of Oculomotor Learning after Changes in Sensory Processing. J Neurosci 37: 11469–11484, 2017. doi: 10.1523/JNEUROSCI.1208-17.2017.

72. Verghese P. Active search for multiple targets is inefficient. Vision Research 74: 61– 71, 2012. doi: 10.1016/j.visres.2012.08.008.

73. Wolf C, Wagner I, Schütz AC. Competition between salience and informational value for saccade adaptation. Journal of Vision 19: 26–26, 2019. doi: 10.1167/19.14.26.

74. Wolf C, Schütz AC. Earlier saccades to task-relevant targets irrespective of relative gain between peripheral and foveal information. Journal of Vision 17: 1–18, 2017. doi: 10.1167/17.6.21.

75. Clarke ADF, Nowakowska A, Hunt AR. Seeing beyond salience and guidance: the role of bias and decision in visual search. Vision (Switzerland) 3: 46, 2019. doi: 10.3390/vision3030046.

76. Gershman SJ, Horvitz EJ, Tenenbaum JB. Computational rationality: A converging paradigm for intelligence in brains, minds, and machines. Science 349: 273–278, 2015. doi: 10.1126/science.aac6076.

77. Rosenholtz R. Demystifying visual awareness: Peripheral encoding plus limited decision complexity resolve the paradox of rich visual experience and curious perceptual failures. Atten Percept Psychophys 82: 901–925, 2020. doi: 10.3758/s13414-019-01968-1.

78. Wise T, Emery K, Radulescu A. Naturalistic reinforcement learning. Trends in Cognitive Sciences 28: 144–158, 2024. doi: 10.1016/j.tics.2023.08.016.

79. Yu X, Zhou Z, Becker SI, Boettcher SEP, Geng JJ. Good-enough attentional guidance. Trends in Cognitive Sciences 27: 391–403, 2023. doi: 10.1016/j.tics.2023.01.007.

80. Shadmehr R, De Xivry JJO, Xu-Wilson M, Shih TY. Temporal discounting of reward and the cost of time in motor control. Journal of Neuroscience 30: 10507– 10516, 2010. doi: 10.1523/JNEUROSCI.1343-10.2010.

81. Shadmehr R. Control of movements and temporal discounting of reward. Current Opinion in Neurobiology 20: 726–730, 2010. doi: 10.1016/j.conb.2010.08.017.

82. Sedaghat-Nejad E, Shadmehr R. The cost of correcting for error during sensorimotor adaptation. Proceedings of the National Academy of Sciences 118: e2101717118, 2021. doi: 10.1073/pnas.2101717118.

83. Callaway F, Van Opheusden B, Gul S, Das P, Krueger PM, Griffiths TL, Lieder F. Rational use of cognitive resources in human planning. Nat Hum Behav 6: 1112–1125, 2022. doi: 10.1038/s41562-022-01332-8.

84. Chukoskie L, Snider J, Mozer MC, Krauzlis RJ, Sejnowski TJ. Learning where to look for a hidden target. Proc Natl Acad Sci USA 110: 10438–10445, 2013. doi: 10.1073/pnas.1301216110.

85. Cohen JD, McClure SM, Yu AJ. Should I stay or should I go? How the human brain manages the trade-off between exploitation and exploration. Phil Trans R Soc B 362: 933–942, 2007. doi: 10.1098/rstb.2007.2098.

86. Gottlieb J, Oudeyer P-Y. Towards a neuroscience of active sampling and curiosity. Nature Reviews Neuroscience 19: 758–770, 2018. doi: 10.1038/s41583-018-0078-0.

87. Johansson RS, Westling G, Bäckström A, Flanagan JR. Eye-hand coordination in object manipulation. Journal of Neuroscience 21: 6917–6932, 2001. doi: 10.1523/jneurosci.21-17-06917.2001.

88. Jóhannesson ÓI, Thornton IM, Smith IJ, Chetverikov A, Kristjánsson Á. Visual Foraging With Fingers and Eye Gaze. i-Perception 7: 204166951663727, 2016. doi: 10.1177/2041669516637279.

89. Moskowitz JB, Fooken J, Castelhano MS, Gallivan JP, Flanagan JR. Visual search for reach targets in actionable space is influenced by movement costs imposed by obstacles. Journal of Vision 23: 4, 2023. doi: 10.1167/jov.23.6.4.

90. Domínguez-Zamora FJ, Marigold DS. Motor cost affects the decision of when to shift gaze for guiding movement. Journal of Neurophysiology 122: 378–388, 2019. doi: 10.1152/jn.00027.2019.

